# Microbial dark matter filling the niche in hypersaline microbial mats

**DOI:** 10.1101/2020.06.18.160598

**Authors:** Hon Lun Wong, Fraser I. MacLeod, Richard Allen White, Pieter T. Visscher, Brendan P. Burns

**Author notes:** Corresponding author: Dr Brendan P. Burns, School of Biotechnology and Biomolecular Sciences, University of New South Wales, Sydney, 2052, Australia.

## Abstract

Shark Bay, Australia, harbours one of the most extensive and diverse systems of living microbial mats, that are proposed to be analogs of some of the earliest ecosystems on Earth. These ecosystems have been shown to possess a substantial abundance of uncultivable microorganisms. These enigmatic groups - ‘microbial dark matter’ (MDM) - are hypothesised to play key roles in microbial mats. We reconstructed 115 metagenome-assembled genomes (MAGs) affiliated to MDM, spanning 42 phyla within the bacterial and archaeal domains. We classified bacterial MDM from the PVC group, FCB group, Microgenomates, Parcubacteria, and Peregrinibacteria, as well as a high proportion of archaeal MDM under the TACK, DPANN, Altiarchaeales, and Asgard archaea. The latter includes the first putative Heimdallarchaeota MAG obtained from any microbial mat system. This study reports novel microorganisms (Zixibacterial order GN15) putatively taking part in dissimilatory sulfate reduction in surface hypersaline settings, as well as novel eukaryote signature proteins in the Asgard archaea. Despite possessing reduced-size genomes, the MDM MAGs are capable of fermenting and degrading organic carbon, suggesting a role in recycling organic carbon. Several forms of RuBisCo were identified, allowing putative CO_2_ incorporation into nucleotide salvaging pathways, which may act as an alternative carbon and phosphorus source. High capacity of hydrogen production was found among Shark Bay MDM. Putative schizorhodopsins were also identified in Parcubacteria, Asgard archaea, DPANN archaea, and Bathyarchaeota, allowing these members to potentially capture light energy. Diversity-generating retroelements were prominent in DPANN archaea that likely facilitate the adaptation to a dynamic, host-dependent lifestyle. In light of our findings, we propose H_2_, ribose and CO/CO_2_ as the main energy currencies of the MDM community in these mat systems.

## INTRODUCTION

A vast ‘known-unknown’ and even ‘unknown-unknown’, many microorganisms have yet to be unlocked from a majority of Earth’s ecosystems. These uncultured microbial community members represent a vast untapped and uncharacterised resource of biological information, representing the ‘microbial dark matter’ (MDM) of many microbial ecosystems [1, 2]. Despite the majority of these unexplored lineages have reduced genome sizes and minimal biosynthetic capacity, they were proposed to represent a major uncharacterised portion of microbial diversity and inhabiting every possible metabolic niche [1-5].

Advances in microbial dark matter research have the potential to alter our understanding of key tenets of evolutionary principles, such as the current debate over the phylogenetic position of the Asgard archaea, and the idea that the eukaryotic cell emerged from within the archaeal domain [6-8]. It is estimated that at least 80% of environmental genomic content could be considered as ‘genomic dark matter’ [5, 9-11], with the majority found in subsurface environments [4, 12-15]. There is indeed a risk that many novel and unexpected organisms and/or their gene products may simply be lost in large datasets. Therefore, this prompted the desire to uncover unknown genes, functions, and ecological roles of these novel groups in other microbial ecosystems [16], of particular interest for the present study, hypersaline microbial mats.

Shark Bay, in Western Australia, harbors one of the most extensive (and diverse) microbial mat systems in the world that are analogs of some of the earliest ecosystems on Earth. Hypersalinity, heatwaves, high UV radiation, oligotrophic waters, fluctuating tides, and even cyclonic events, contribute to the mats being subjected to an extreme environment in Shark Bay [17-21]. Indeed recent findings have shown that these mats are subjected to extreme fluctuations over a diel cycle as tides change [22-24], both at the surrounding microenvironment level (large changes in temperature, salinity, pH) as well as at the metabolic level (rapid changes in O and S levels reflective of changing rates of photosynthesis, respiration, and sulfur metabolisms). Despite recent advances made in our understanding of Shark Bay mat taxonomic and functional complexity [17, 19, 20, 22-24], the diversity and ecological role of MDM in these evolutionarily significant systems is unknown. Amplicon sequencing revealed that MDM comprises over 15% of the bacterial community and over half of the archaeal population in Shark Bay mats [22, 23]. Although these results have indicated Shark Bay microbial mats are a huge genetic pool of novel lineages, their functional role(s), including how they adapt to such an extreme environment and their putative interactions with other microorganisms, are still unknown.

We hypothesize that microbial dark matter in microbial mats may be key in nutrient cycling, symbioses, and overall health of these systems under extreme environmental conditions. In this study we have unravelled in detail the metabolic potential and capacities of this enigmatic group of novel microorganisms in Shark Bay mats.

## RESULTS AND DISCUSSION

### Microbial dark matter metagenome-assembled genomes (MAGs)

This study describes for the first time in detail MAGs associated with microbial dark matter in hypersaline microbial mats. In total, 115 MAGs were found in Shark Bay mat metagenomic data [24], spanning 42 phyla within the bacterial and archaeal domains (Fig. 1). MAGs that were classified as part of microbial dark matter in previous literature were chosen in this study [1, 3-6, 12, 13, 25, 26]. Genome statistics are provided in Additional file 13: Table S1. Of the 115 MAGs, 24 high-quality MAGs were obtained (> 90% completeness, < 5% contamination, encoding at least 18 out of 20 amino acids) and the remaining were of medium quality MAGs (> 50% completeness, < 10% contamination) based on recently established standards [27]. Although one Heimdallarchaeota (Bin_120) and one Lokiarchaeota (Bin_186) MAG had slightly over 10% contamination levels (both with 10.75%), they are included in this study due to high completeness (> 85%), as well as this being the first Heimdallarchaeota MAG obtained from any microbial mat system. Of the 91 medium quality MAGs, 65 have > 70% completeness. Bacterial MAGs were further classified into the PVC group, FCB group, Microgenomates, Parcubacteria, Peregrinibacteria, and ‘others’ (Fig. 1). Archaeal MAGs were classified into Asgard archaea, TACK, DPANN, and Altiarchaeales. It is still under debate whether Altiarchaeales should be placed within the DPANN superphylum [28-30], so these were placed as a separate group in the current analyses.

**Figure 1.**
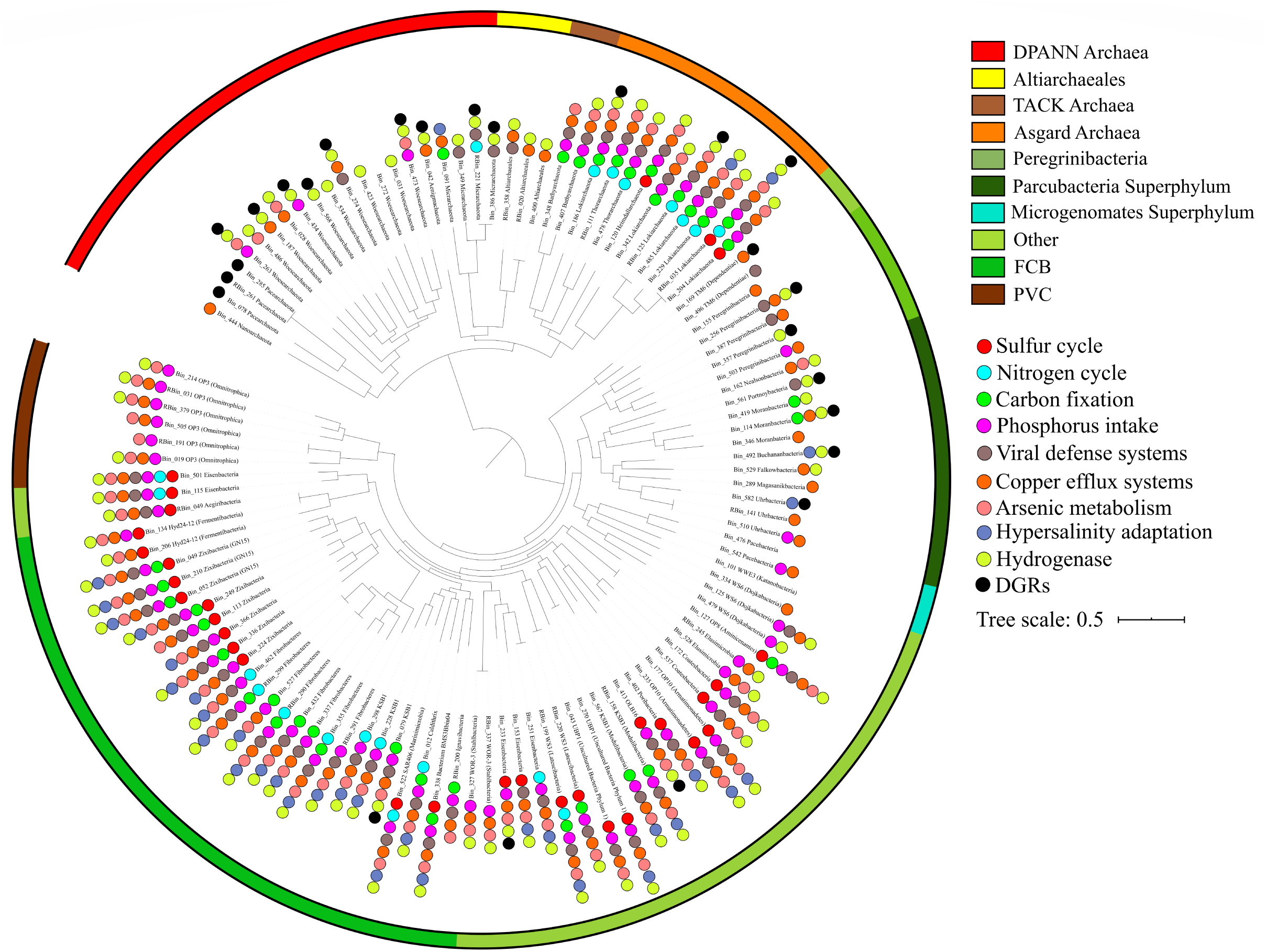
Phylogenetic tree of novel MAGs of the MDM in Shark Bay microbial mats. Maximum likelihood phylogenetic tree of up to 16 concatenated ribosomal proteins (rpL2, 3, 4, 5, 6, 14, 15, 16, 18, 22, 24 and rpS3, 8, 10, 17, 19) was constructed. Bin_245 (Bathyarchaeota) is not included in the tree as it has less than 8 ribosomal proteins. MAGs assigned to different groups are highlighted in different colors on the outer circular stripe. Circles represent genes for various nutrient cycles present in the MAGs (detailed in Additional file 15: Table S3).

### Distribution of novel rhodopsins and eukaryotic signature proteins (ESPs)

Rhodopsin genes were identified in Asgard archaea, Parcubacteria, Bathyarchaeota, and DPANN archaeal MAGs. Phylogenetic analysis of the rhodopsins showed affiliation with schizorhodopsins, which was recently found in Asgard archaea in a microoxic niche setting [31] (Additional file 2: Figure S1 and Additional file 14: Table S2). This study expanded the range of phyla encoding schizorhodopsin, which has not been identified in Bathyarchaeota, DPANN archaea, and Parcubacteria previously. The novel rhodopsin may infer light sensitivity in these microbial groups with a recent study indicating Asgard archaeal schizorhodopsins as light-driven H^+^ pumps [32]. Rhodopsins in MDM (Saccharibacteria; Asgard archaea) have only been found in hypersaline environments to date, suggesting the acquisition of such an evolutionary adaptation of this enigmatic group may be a feature in hypersaline, sunlit surface settings, with the ability to utilize light energy to counter the high cost of osmotic balance maintenance [31, 33, 34].

Eukaryotic signature proteins (ESPs) were found distributed in the ten Asgard archaeal MAGs (Additional file 1: Supplementary Information and Additional file 3: Figure S2), hinting a close evolutionary relationship between Asgard archaea and eukaryotes [7, 8, 32, 35]. ESPs in Shark Bay Asgard archaeal MAGs are putatively involved in cytoskeleton dynamics, information processing, trafficking machinery, signalling systems and N-linked glycosylation [7, 8]. Novel ESPs were identified affiliated with information processing and GTP binding proteins belonging to ARF, RAS, RAB and RAN families in Asgard archaea for the first time, suggesting that this archaeal superphylum possess a diverse range of eukaryotic-like signalling systems (Additional file 3: Figure S2).

### Limited metabolic capacities and putative host-dependent lifestyle

Most of the MDM MAGs harbor incomplete metabolic pathways despite the oligotrophic condition present in Shark Bay (Additional file 1: Supplementary Information), hence are suggested to be host-dependent [29, 36]. We screened the MAGs for diversity-generating retroelements (DGRs), which are fast-evolving proteins enabling host-dependent microorganisms to attach to the hosts’ surface [30, 37-39], and they were identified mostly in Parcubacteria and DPANN archaea (Fig. 1 and Fig. 2). Interestingly, despite having versatile metabolic pathways, DGRs were also identified in Asgard archaeal MAGs in this study, which has not been reported before (Fig. 1, Fig. 2 and Additional file 1: Supplementary Information). This may indicate Asgard archaea once resided in energy-limited environments which relies on symbiotic relationships [32, 37]. Parcubacteria and DPANN archaeal members lack biosynthetic capabilities and were suggested to have a symbiotic, host-dependent lifestyle, in which DGRs facilitate adaptation to rapidly changing environments, providing them with the tools to exploit symbiotic associations with their host [1, 29, 30, 40].

**Figure 2.**
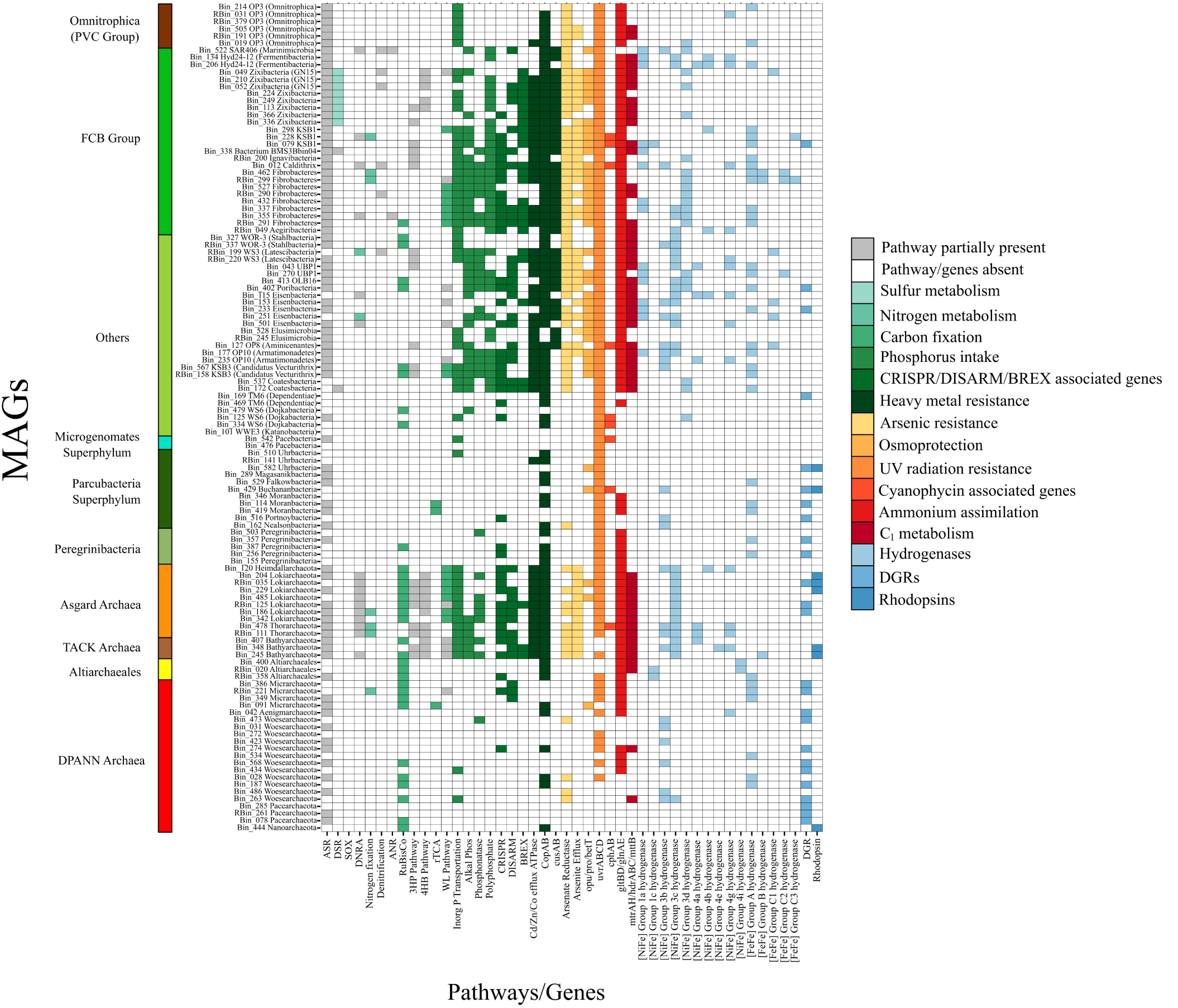
Color-coded table indicating major functional genes identified in Shark Bay mat novel microbiome MAGs. X-axis indicates specific genes likely involved in either nutrient cycling or environmental adaptation and y-axis indicates the specific microbial dark matter MAGs. Key: Grey indicates the partial pathways identified in carbon, sulfur and nitrogen cycles; white indicates the genes and associated pathways are absent. Color panel on the left represents different groups of MDM MAGs according to Figure 1. ASR, assimilatory sulfate reduction; DSR, dissimilatory sulfate reduction; SOX, sulfur oxidation; DNRA, dissimilatory nitrate reduction; ANR, assimilatory nitrate reduction; rTCA, reverse tricarboxylic cycle; WL pathway, Wood-Ljungdahl pathway; Inorg P, inorganic phosphorus; Alkal Phos, alkaline phosphatase; copAB/cusAB, copper efflux systems; opu, osmoprotectant transport system; pro, glycine betaine/proline transport system; bet, choline/glycine/proline betaine transport protein; uvr, exinuclease; cph, cyanophycin; gltBD/glnAE, ammonium assimilation; mtr, tetrahydromethanopterin S-methyltransferase; hdr, heterodisulfide reductase; mttB, trimethylamine-corrinoid protein co-methyltransferase; DGR, diversity-generating retroelements.

It is hypothesised that MDM may take advantage of the lack of virus defence systems observed here (CRISPR, BREX, DISARM and DNA phosphorothioation; Additional file 1: Supplementary Information and Additional file 15: Table S3), despite the high abundance of viruses and virus defence systems identified in Shark Bay in previous studies [24, 41]. Given the limited metabolic capacities and host-dependent lifestyle, these organisms may serve as ‘viral decoys’ to alleviate the load on the host’s viral defence system, and benefits from the viral DNA as a pentose and phosphorus source [42]. Another advantage is to avoid autoimmunity, in which the virus defence systems may attack the host [42, 43]. Furthermore, maintaining such systems is energetically costly [44]. It is proposed that the synergy between the lack of viral defence systems and the presence of DGRs facilitate rapid screening and acquisition of biological functions for survival in these mat systems, allowing MDM with limited metabolic pathways to adapt to a dynamic and extreme environment.

### Sulfur, nitrogen, and carbon cycling in Shark Bay MDM

Despite having reduced metabolic capacity, various genes within sulfur, nitrogen, and carbon cycles were found distributed among the Shark Bay microbial dark matter community. All Zixibacteria and Zixibacteria order GN15 (previously classified as candidatus phylum GN15) genomes harbour dissimilatory (bi)sulfite reductase (*dsrAB*) and adenylylsulfate reductase (*aprAB*) genes affiliated with dissimilatory sulfate reduction [45, 46] (Additional file 4: Figure S3 and Additional file 15: Table S3). MAGs of unclassified bacterium BMS3Bbin04 (FCB Group) also harbour *dsrAB*, while Coatesbacteria encode for an *aprA* gene, suggesting these bacteria may play a role in the sulfur cycle. To confirm their role(s) in the sulfur cycle, *dsrAB* genes were analysed against the *dsrAB* database [47]. The genes were classified as a reductive bacterial type, confirming their likely function as dissimilatory (bi)sulfite reductase (Additional file 5: Figure S4). The *dsrAB* genes clustered with uncultured lineages in estuarine environments and interestingly, one arctic *dsrAB* lineage, suggesting a specific lineage in Shark Bay (Additional file 5: Figure S4). Furthermore, *dsrC* genes were found co-localised on the same contigs in the *dsrAB* encoding MAGs (Additional file 15: Table S3), which is an essential physiological partner to *dsrAB* in sulfite reduction [46]. This suggests Zixibacteria (and order GN15) potentially partake in dissimilatory sulfate reduction not only in deep subsurface [46, 48, 49], but also in surface hypersaline environments. Genes *dsrEFH* were also identified in 40 MAGs in the present study, expanding the lineages taking part in sulfur cycling (Additional file 1: Supplementary Information). Previous studies in these mat systems have indicated a putative surface anoxic niche with high rates of sulfur cycling [22, 24], and potentially microbial dark matter could be contributing to this phenomenon.

Only a limited number of genes involved in nitrogen cycling were identified in MDM (Fig. 1, Fig. 3 and Additional file 15: Table S3). Genes encoding nitrite reductase were found in all Asgard archaea MAGs except Heimdallarchaeota (Fig. 2 and Additional file 15: Table S3). The co-occurrence of CO dehydrogenase and nitrite reductase suggests that Asgard archaea may potentially couple CO oxidation to nitrite reduction [50], allowing them to derive energy from an oligotrophic environment (Fig. 2, Fig. 4 and Additional file 15: Table S3).

**Figure 3.**
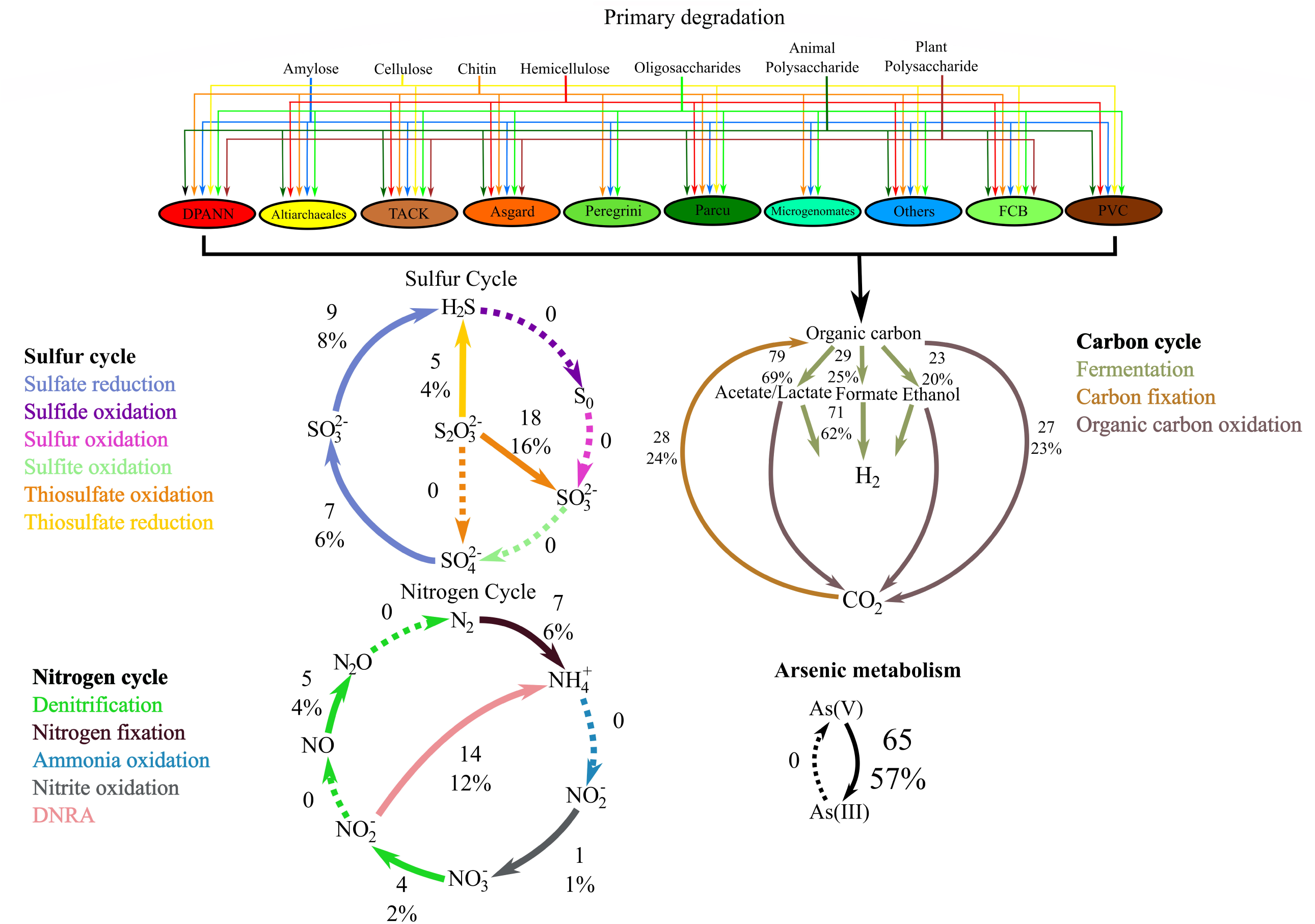
Putative involvement of Shark Bay MDM MAGs in carbon, sulfur, nitrogen and arsenic metabolisms. CAZy enzymes with different colored arrows representing various groups of glycoside hydrolase corresponding to Additional file 6: Figure S5. Numbers indicate the quantity and percentage of MAGs encoding for the nitrogen/sulfur cycles and metabolic pathways.

**Figure 4.**
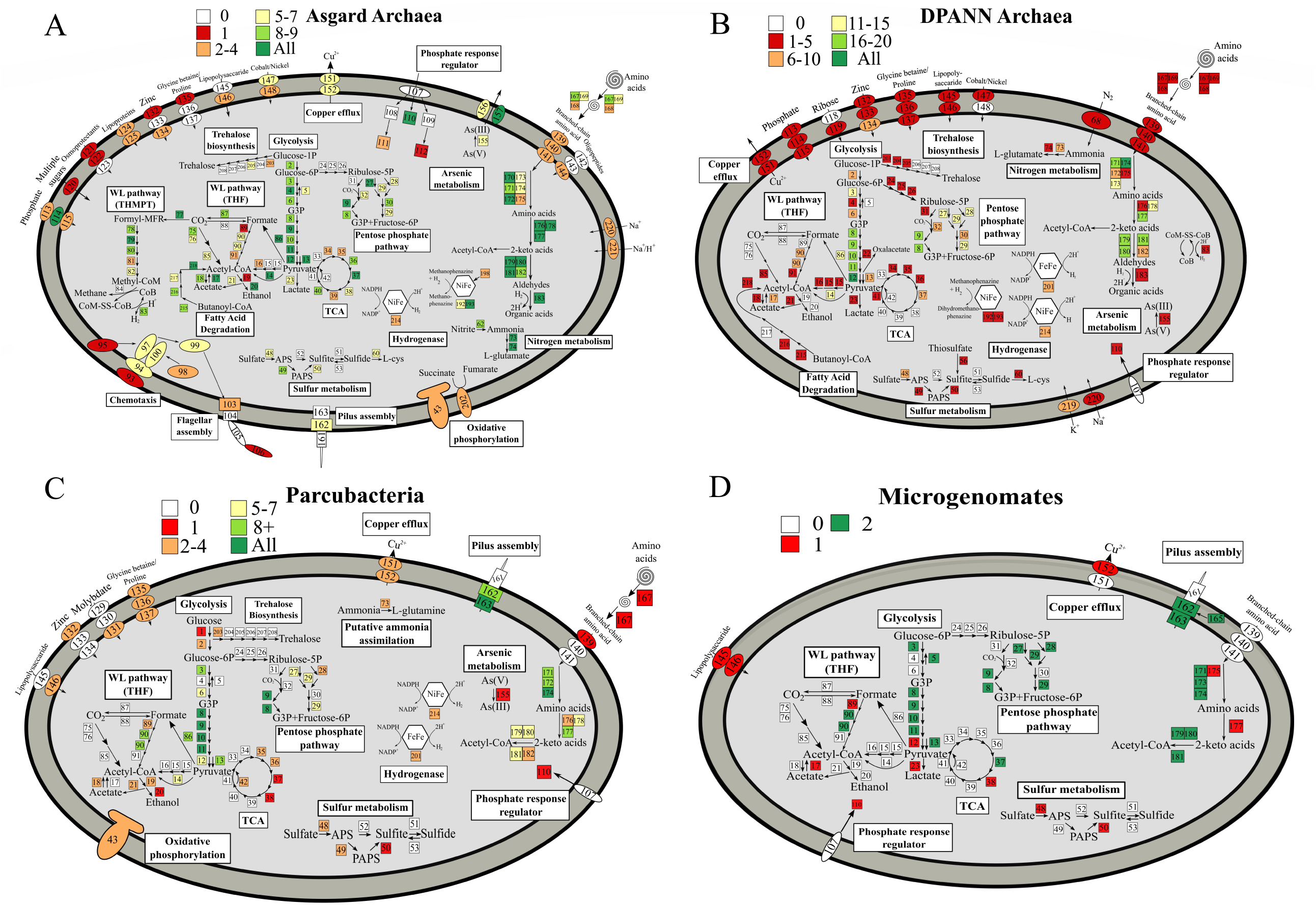
Metabolic potential of Asgard archaea, DPANN archaea, Parcubacteria and Microgenomates in the Shark Bay systems. A metabolic map summarising the genomic potential and metabolic capacities of MAGs affiliated with: **A.** Asgard archaea MAGs **B.** DPANN archaea MAGs **C.** Parcubacteria MAGs **D.** Microgenomates MAGs. Numbers represent specific genes in given pathways and the corresponding genes are listed in Additional file 15: Table S3. Different colors in the square boxes represent different numbers of MAGs encoding the genes, while white square boxes indicate the absence of the genes. TCA, tricarboxylic acid cycle; THF, tetrahydrofolate; THMPT, tetrahydromethanopterin; WL pathway, Wood-Ljungdahl pathway; PAPS, 3’-phosphoadenylyl sulfate; APS, Adenylyl sulfate.

To infer the capacity of carbohydrate degradation in microbial mat microbial dark matter, we analysed MAGs for carbohydrate-active enzymes (CAZy). Asgard archaea and MAGs affiliated within the FCB group have the broadest cassette of glycoside hydrolase (GH) genes, (hemicellulose, amylase, animal and plant polysaccharides), indicating a highly flexible metabolic capacity for carbon acquisition (Fig. 3, Additional file 1: Supplementary Information and Additional file 6: Figure S5). On the other hand, Parcubacteria, Microgenomates, Peregrinibacteria, and DPANN archaea encode a lower range of GH enzymes, suggesting these members could scavenge readily degraded carbohydrates through their potential symbiotic hosts or partners. Most of the microorganisms identified in this study are likely capable of fermenting various carbon sources into formate, acetate, lactate, and ethanol (Fig. 3, Fig. 4 and Additional file 15: Table S3). This finding suggests that most MDM MAGs undergo anoxic carbon transformation, corroborating with previous studies [13, 14, 40, 51-53].

Genes encoding anaerobic carbon monoxide dehydrogenase (*cooSF*) and acetyl-CoA synthase (*cdhDE, acsB*) were identified in FCB group MAGs (Modulibacteria, KSB1, Fibrobacteres) (Additional file 4: Figure S3), Asgard archaea (Heimdall-, Loki-, Thorarchaeota) (Fig. 4) and Bathyarchaeota (Additional file 7: Figure S6) MAGs, indicating their putative ability to fix and reduce CO_2_ to acetyl-CoA through the Wood-Ljundahl (WL) pathway (Fig. 2 and Additional file 15: Table S3). Generally, microorganisms can either use tetrahydrofolate (THF) or tetrahydromethanopterin (THMPT) as C_1_ carriers in the WL pathway [54, 55]. Usually, bacteria utilise the THF-WL pathway while THMPT-WL pathways are mostly found in archaea [54-57]. One Lokiarchaeota MAG (Bin_186) harbours a complete anaerobic H_2_-dependent THMPT-WL pathway, inferring the ability to fix CO_2_ and H_2_ into acetate [58] (Fig. 4 and Additional file 15: Table S3). All other Asgard archaea in the Shark Bay mats contained most genes for both WL pathways, though incomplete genomes may account for the absence of these genes (Additional file 15: Table S3).

The presence of the Wood-Ljungdahl pathway is suggested to be a result of energy limitation since it is energetically inexpensive compared to other carbon fixation pathways [45, 58, 59]. Up to a quarter of the MAGs in the present study encode for CO dehydrogenase, allowing CO to be putatively utilised [60, 61]. Given the high UV radiation the Shark Bay mats are exposed to, CO may be produced through photo-degradation and subsequently oxidised as an alternative carbon source for energy conservation [24, 62, 63].

Furthermore, based on the observed genomic repertoires, Asgard archaea in these mats are putatively heterotrophic acetogens, encode for a complete beta-oxidation pathway and may take part in the carbon fixing 4-hydroxybutyrate pathway (Additional file 1: Supplementary Information). The MDM community has scattered genes in other carbon metabolisms but encode peptidases, putatively facilitating scavenging organic carbon in their oligotrophic environment (Additional file 1: Supplementary Information).

### Nucleotide salvaging and putative CO_2_ assimilation

Surprisingly, despite the reduced-sized genomes, 32 MAGs encode for ribulose biphosphate carboxylase (RuBisCo) (Fig. 2, Additional file 15: Table S3 and Additional file 16: Table S4). Given not all types of ribulose biphosphate carboxylase undergo carbon fixation, a phylogenetic tree was constructed to examine the variety of RuBisCo in these mat metagenomes. The MDM MAGs appear to harbour bacterial and archaeal type III, type IIIa, type IIIb, type IIIc, and type IV RuBisCo (Fig. 5). Furthermore, all MAGs have incomplete CBB (Calvin-Benson-Bassham) cycle (Additional file 15: Table S3). This suggests that these microorganisms are involved in the AMP nucleotide salvaging pathway, while MAGs harbouring type IV RuBisCo are involved in methionine salvage pathways [64, 65]. Interestingly, 22 out of the 32 MAGs with RuBisCo also encode both AMP phosphorylase (*deoA*) and R15P isomerase (*e2b2*) (Additional file 15: Table S3), indicating the potential ability to incorporate CO_2_ into nucleotide salvaging pathways [52, 65-67]. One Lokiarchaeota MAG (Bin_186) encodes for a type IIIa RuBisCo, which is known to fix CO_2_ through the reductive hexulose-phosphate (RHP) cycle [65, 66]. Although it potentially lacks the ability to fix CO_2_ due to the absence of homologs to genes encoding phosphoribulokinase, this MAG encodes for a fused bifunctional enzyme 3-hexulose-6-phosphate synthase/formaldehyde-activating enzyme (*fae-hps*) allowing for the potential production of methylene-H_4_MPT, which may play a role in replenishing the C_1_ carriers in the THMPT-WL pathway [66].

**Figure 5.**
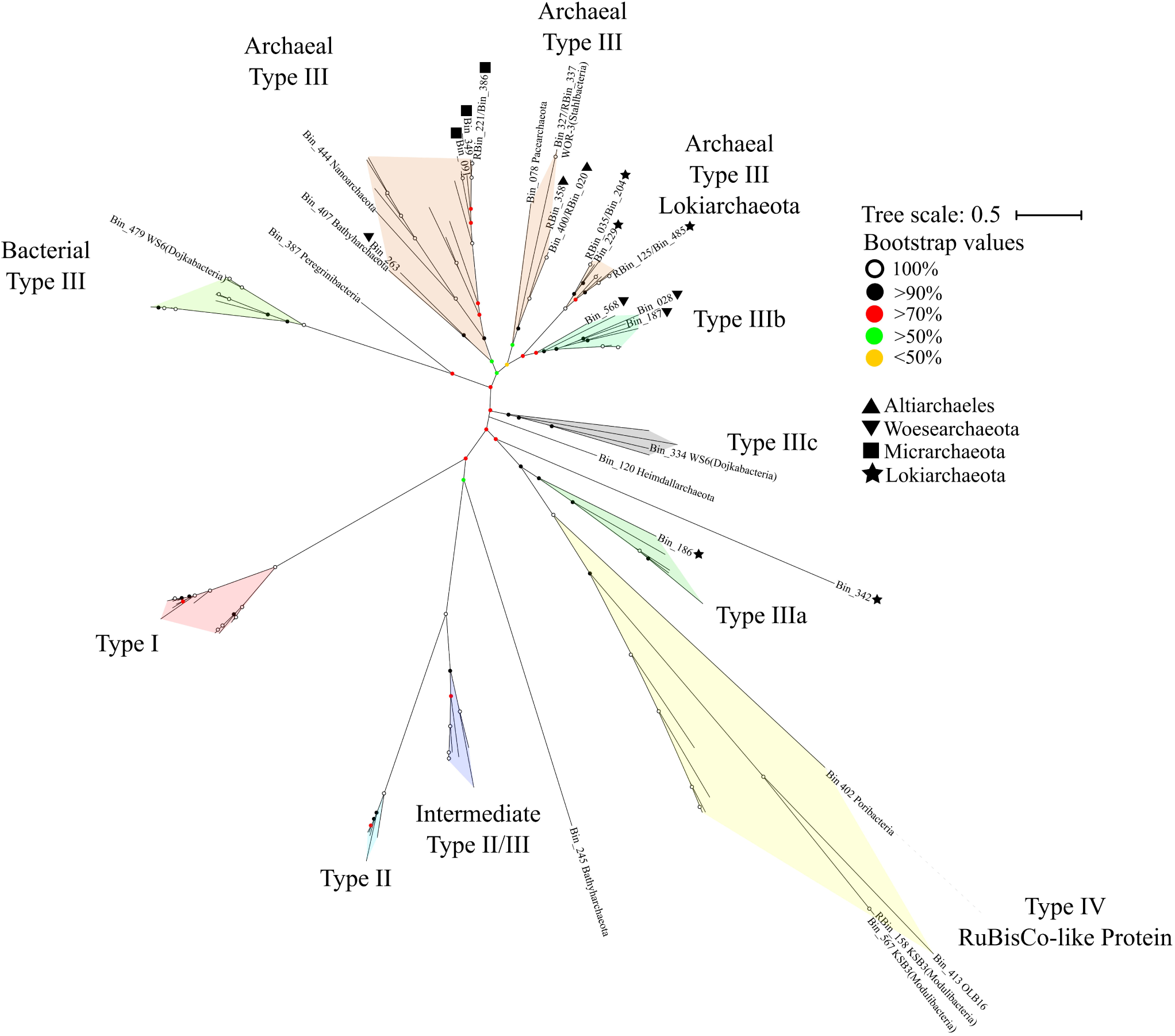
Unrooted maximum-likelihood phylogenetic tree of RuBisCo genes in Shark Bay MDM MAGs. Maximum-likelihood phylogenetic tree constructed with RuBisCo gene found in the MDM MAGs with 1000 bootstrap replications. Archaeal and bacterial type III, type IIIa [66, 67], type IIIb [46], type IIIc [46] and type IV RuBisCo-like protein [64] were identified. Circular dots of different colors represent bootstrap values. RuBisCo sequences in this study and reference sequences are listed in Additional file 16: Table S4.

One interesting finding is that Heimdallarchaeota (Bin_120) contains RuBisCo at the basal position (Fig. 5), suggesting it may possess RuBisCo as an early-evolved form. The widespread distribution of RuBisCo among MDM in Shark Bay mats implies the use of ribose to substitute upper glycolysis, as some of the key genes in this pathway are missing [52] (Additional file 1: Supplementary Information). Other than feeding ribose (and putatively CO_2_) as augmented carbon sources into the central carbon metabolism, these non-autotrophic RuBisCo may putatively free phosphate groups from nucleotides to supplement the extremely limited phosphorus in Shark Bay found in previous studies [24, 68]. In this study the genomes harbouring RuBisCo were identified across 14 MDM phyla, suggesting that ribose may be a prominent currency among microbial dark matter in hypersaline microbial mats (Fig. 4, Fig. 5, Additional file 7-11: Figure S6-S10).

### High capacity for hydrogen production among Shark Bay MDM

A total of 267 hydrogenases were detected in 81 out of 115 MDM MAGs, implying prominent hydrogen metabolism in MDM even with minimal genomes (Fig. 4, Additional file 4: Figure S3, Additional file 7-11: Figure S6-10 and Additional file 15: Table S3). A total of 16 types of hydrogenases were identified against the HydDB database [69], including 10 [NiFe] and 6 [FeFe] classes (Fig. 2, Additional file 15: Table S3 and Additional file 1: Supplementary Information). Most of the hydrogenases identified are putatively involved in H_2_ uptake (Group 1), H_2_ consumption/production, fermentative H_2_-evolving, and H_2_ sensing (Group 3, 4 and [FeFe]) (Fig. 2).

Almost half of the Shark Bay MAGs harbour hydrogenases associated with H_2_ production, and of particular significance, Parcubacteria and Woesearchaeota only encode H_2_ producing hydrogenases ([NiFe]-3b and [FeFe] Group A), which are fermentative in nature [70] (Fig. 2, Fig. 4 and Additional file 15: Table S3). H_2_ production is potentially an important energy currency in these mats as hydrogenotrophic methanogenesis was found to be the prominent mode of methane production [23]. Furthermore, a global survey suggests that Woesearchaeota form consortiums with hydrogenotrophic methanogens by providing H_2_ in exchange of nutrients [71]. It is suggested that these MAGs (especially among Parcubacteria and Woesearchaeota) support and complement H_2_/CO_2_ methanogenesis in Shark Bay microbial mats.

One-third of the MAGs (43 out of 115) encodes for 3b and 3c hydrogenases, which play essential roles as electron donors and H_2_ production during hydrogenogenic fermentation and Wolfe cycle of methanogenesis [72, 73]. Of particular interest, the presence of the WL pathway along with hydrogenase group 3b and 3c in Asgard archaea suggests this group are putatively lithoautotrophs that use H_2_ as electron donors [58, 74]. With a range of CAZy enzymes distributed among Shark Bay MDM (Additional file 6: Figure S5), these microorganisms likely participate in anoxic carbon transformations and hydrogen turnover [12, 14, 40, 75-77]. Therefore, MDM in these systems may act as a ‘recycler’ in the mats to recycle organic carbon from dead cells, employing hydrogenogenic or hydrogenotrophic metabolisms.

### Energy Currencies of MDM

As described in an earlier study [24], it is likely that the WL-pathway is the main mode of carbon fixation in these mats, and the surface phototrophic consortia produce the energy and organic carbon for the rest of the microbial community [22, 78]. Various adaptation strategies to the hypersalinity, limited phosphorus, and high copper concentration were described [24] (Additional file 1: Supplementary Information). However, given the oligotrophic nature of Shark Bay waters [79], MDM in these mats that lacks the metabolic capacity may utilise alternative carbon sources to augment nutrient intake. First, it is proposed that due to the high UV irradiation in Shark Bay, photo-degradation of surface organic matter may provide CO as an alternative carbon source [62, 63]. Secondly, the widespread hydrogenases among MDM may contribute to the hydrogen turnover in exchange of nutrients as high rates of hydrogenotrophic methanogenesis were measured and detected in these mats [23]. Thirdly, RuBisCo found in the MDM MAGs is proposed to fix CO_2_ alongside nucleotide salvaging, which is subsequently fed into glycolysis, maximising energy yield [52, 66, 80, 81]. It is therefore proposed in an ecological context, MDM occupies metabolic niches in Shark Bay microbial mats where ribose, H_2_, CO and CO_2_ are prominent currencies to augment energy income.

## Conclusions

This is the first study to reconstruct and describe in detail high-quality genomes affiliated with microbial dark matter in microbial mats. This study reports the novel uncultured bacterial phyla Zixibacteria (including order GN15) and an unidentified bacterium (Bin_338) as likely participants in dissimilatory sulfate reduction in surface hypersaline settings, as well as diversity generating retroelements and novel ESPs identified in Asgard archaea. It is suggested that Asgard archaea are not only organoheterotrophs, but also putatively lithoautotrophs that have more versatile metabolic capacities than the other groups of MDM, possessing both THMPT- and THF-WL pathways, RuBisCo and schizorhodopsin. For the other MDM groups, although possessing minimal genomes and the lack of complete biosynthetic pathways, they are potentially capable of degrading and fermenting organic carbon and are suggested to play a role in H_2_ and carbon transformation in microbial mats. Various forms of RuBisCo were encoded, allowing putative CO_2_ incorporation into nucleotide salvaging pathways, acting as an alternative carbon and phosphorus source. Despite possessing minimal genomes, DGRs were prominent in Parcubacteria and DPANN archaea to likely adapt to a dynamic, host-dependent environment. Under the oligotrophic environment in Shark Bay, MDM needs to exploit every opportunity for energy generation, such as harbouring scattered genes of various nutrient cycles to fill in metabolic gaps or function in “filling the niches” and as is the case for some other ecological systems [12, 30, 82] (Additional file 1: Supplementary Information). On the other hand, MDM in Shark Bay may be shaping the mat environment through their various metabolic capacities in a process called niche construction, modifying their own and each other’s niches and functional roles in the ecosystem [16, 83, 84]. It has in fact been recently suggested that early ecosystems such as microbial mats were not nutrient starved but rather limited by electron donor/acceptor availability [85], thus the ability for these ecosystems to maximize energy yielding capacities is evolutionary advantageous. A conceptual ecological model of MDM in Shark Bay mats is shown in Fig. 6, proposing the MDM serves to fill in the metabolic gaps. Ribose, CO_2_/CO, and H_2_ are suggested to be prominent currencies among MDM in these mats and were potentially a widespread phenomenon on early Earth [40, 86].

**Figure 6.**
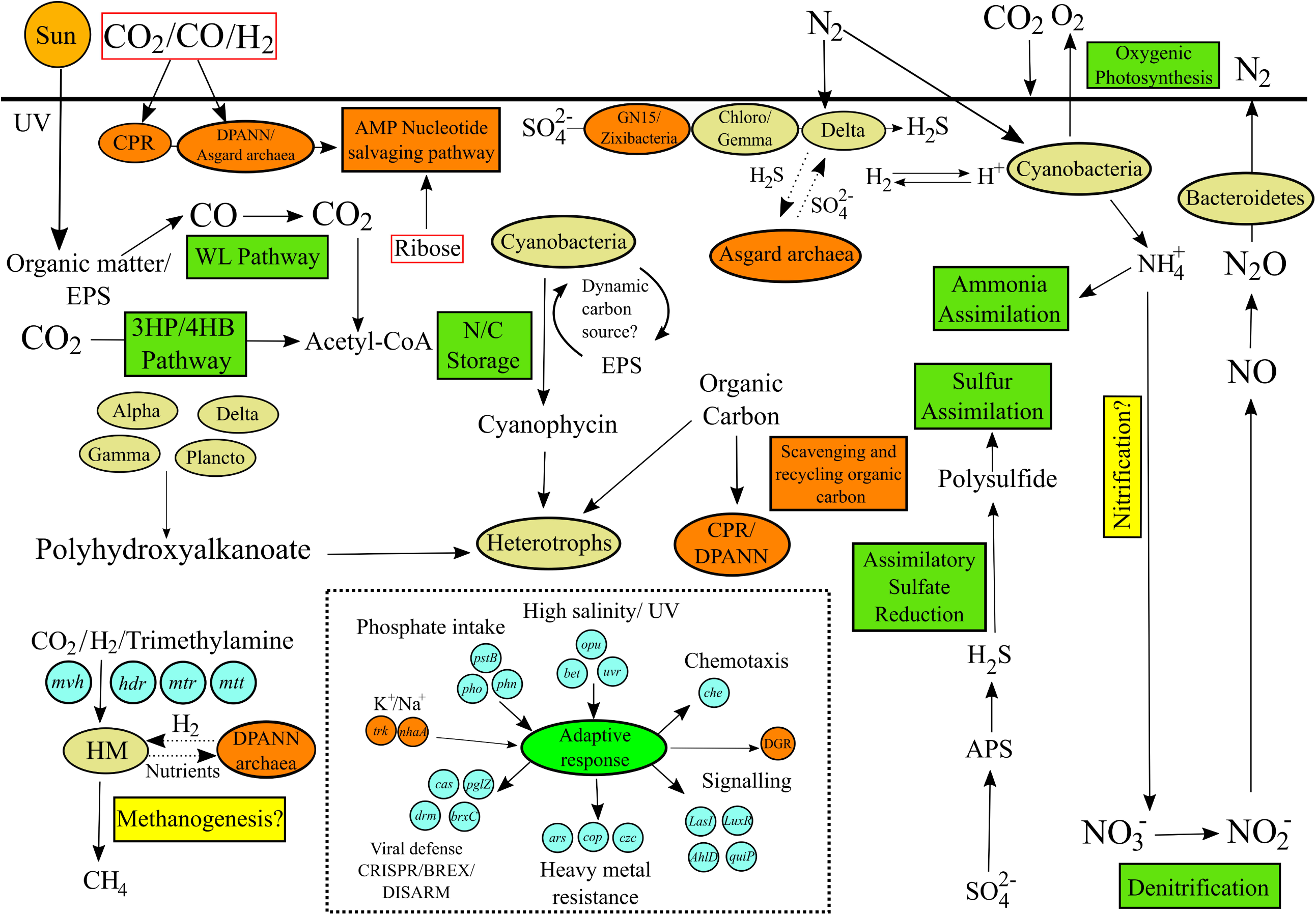
Proposed ecological model of Shark Bay mats (modified from Wong et al., 2018 [24], incorporating MDM). Green rectangular boxes indicate metabolic pathways, yellow rectangular boxes indicate putative pathway and orange rectangular boxes represent putative functional roles of MDM in these mats. Dark green ovals represent microorganisms and their corresponding functions [24] while orange ovals represent microbial dark matter. Red boxes encircle CO_2_/CO, H_2_ and ribose that are proposed as main energy currencies of Shark Bay MDM. Dashed arrows indicate putative metabolic exchange/microbial interactions. Dashed box includes genes involved in environmental adaptation. Chloro, Chloroflexi; Gemma, Gemmatimonadetes; HM, hydrogenotrophic methanogens; N/C storage, nitrogen/carbon storage; WL pathway, Wood-Ljungdahl pathway; 3HP/4HB pathway, 3-hydroxypropionate/4-hydroxybutyrate pathway; EPS, extracellular polymeric substance; DGR, diversity-generating retroelements.

## MATERIALS AND METHODS

### Sampling and metagenomic sequencing

Microbial mat sampling from Shark Bay was performed in a previous study [22], and DNA extraction and sequencing of total community DNA are described previously [24]. Metagenomes were analysed from smooth mats in the present study. The Fastq sequencing data files obtained from the Illumina NextSeq platform detailed by Wong and colleagues [24] were used in the present study for detailed analysis of microbial dark matter.

### Assembly, binning and phylogenetic analyses

Low-quality bases (per base sequence quality < 28) from each sequencing file were trimmed and examined using Trimmomatic (version 0.36) and FastQC (version 0.11.6) respectively [87, 88]. All sequencing files from the ten layers of smooth mats were co-assembled as described [89] (minimum kmer 27, with incremental kmer set as 10) using Megahit version 1.1.1 [90]. Subsequently, all contigs with length less than 2000 bp were removed to avoid ambiguous contig annotation of shorter contigs. This step is to avoid misinterpretation of novel annotated contigs. An alignment algorithm, BWA-MEM (version 0.7.7), was used to map reads back to the assembled contigs [91]. SAMtools version 1.3.1 was used to convert SAM files to binary format BAM files [92]. MetaBAT2 (Version 2.12.1), MaxBin2 (Version 2.2.3) and CONCOCT (Version 1.0.0) were applied for metagenomic binning [93-95]. Subsequently, DAS Tool was used to refine and filter lower quality MAGs generated from the three binning programs [96]. CheckM was employed to examine the quality (completeness and contamination level of MAGs), and MAGs statistics were obtained through QUAST and tRNAScan-SE [97-99]. MAGs with at least medium quality (>50% completeness, <10% contamination) were selected in this study [27]. Subsequently, the taxonomy of MAGs was determined with GtDb-tk [100], in which only MAGs classified as microbial dark matter in previous literatures were chosen [1, 3-6, 12, 13, 25, 26].

Maximum-likelihood-based phylogenetic trees based on 16 concatenated ribosomal proteins (rpL2, 3, 4, 5, 6, 14, 15, 16, 18, 22, 24 and rpS3, 8, 10, 17, 19) were constructed as described in Hug et al (2013) [101]. Only MAGs with at least 8 ribosomal proteins were included in the analysis. Bin_245 (Bathyarchaeota) was not included in the phylogenetic tree (Fig. 1) as it has less than 8 ribosomal proteins. Phylosift version 1.0.1 was used to extract ribosomal proteins from the genomes [102]. Subsequently, ribosomal protein sequences were aligned using MAFFT version 7.310 [103]. BMGE was then used to remove gaps in the alignment with BLOSUM30 matrix and gap rate cut-off of 50% [104]. The resulting protein alignments were concatenated as described in Hug et al (2013) [101]. The concatenated ribosomal proteins were then used to construct a phylogenetic tree using IQ-TREE version 1.6.1 with a total of 1000 bootstrap replicates, which the output file was visualised with iTOL [105, 106].

### Functional annotation

Nucleotide contigs of metagenome-assembled genomes were translated to amino acid sequences by employing Prodigal version 2.6.3 [107]. Functional annotation was carried out using GhostKoala to assign amino acid contigs to MAGs against the KEGG database [108]. InterProScan version 5.25-64.0 was employed to annotate protein domains of MAGs to PFAM and TIGRfam databases, with cutoff value <1e^-10^ [109]. Contigs were annotated against the CAZy database to identify carbohydrate-active enzymes in the MAGs [75]. DGRScan was used to identify diversity-generating retroelements (DGR) among the MAGs [110]. Hydrogenase sequences derived from KEGG and PFAM databases were extracted and annotated against HydDB to further classify hydrogenases [69]. ESP and rhodopsin sequences were submitted to HHPred [111] to confirm their identity.

### Phylogenetic analysis of RuBisCo, Rhodopsin and *dsrAB*

To determine the type of RuBisCo identified in MAGs presented in this study, RuBisCo sequences were downloaded from NCBI and ggkBase [52] (http://ggkbase.berkeley.edu). Reference sequences of RuBisCo and rhodopsins are listed in Additional file 16: Table S4 and Additional file 14: Table S2. Reference sequences of dissimilatory sulfate reduction *dsrAB* were obtained from the dsrAB reference database [47]. RuBisCo, rhodopsin, and *dsrAB* sequences were aligned with MAFFT version 7.310 [103], with gaps subsequently removed by UGENE [112]. IQ-TREE version 1.6.1 was employed to construct a phylogenetic tree with a total of 1000 bootstrap replicates and visualised with iTOL [105, 106]. To further confirm the identity of RuBisCo and rhodopsins, the sequences were annotated against the HHpred [111] and the BLAST database [113].

## ACKNOWLEDGEMENTS

The authors would like to thank the Ramaciotti Centre for Functional Genomics for valuable technical support. This research received no specific grant from any funding agency in the public, commercial, or not-for-profit sectors.

