## Supplementary Information for "Microbial dark matter filling the niche in hypersaline microbial mats"

Additional file 1: **Supplementary information.** Supplementary text with additional information.

**Supplementary information accompanying Wong et al (*Microbial dark matter filling the niche in hypersaline microbial mats*)**

**Overall taxonomic contribution of microbial dark matter (MDM) to Shark Bay microbial communities**. Bacterial and archaeal 16S rRNA genes were obtained from Wong et al (2015) [1] and Wong et al (2017) [2] respectively. MOTHUR version 1.33.0 [3] was used to classify OTUs as described in previous studies [1, 2]. Samples were subsampled to 50,000 sequences and were classified against SILVA database Version 132 [4] to obtain 16S rRNA data affiliated to microbial dark matter. Smooth mats have over 13% relative abundance of bacterial MDM (Additional file 17: Table S5), with Woesearchaeota the dominant archaeal phylum, occupying 38.5% of the archaeal population (Additional file 18: Table S6). Asgard archaea comprise 10% of the archaeal 16S rRNA gene sequences, implying a more diverse community of archaeal dark matter in these systems than previously thought. Although most of the novel phylum comprises less than 0.1% of the total bacterial population (Additional file 17: Table S5), it demonstrates the ability of metagenomics in reconstructing genomes affiliated to the uncultured biosphere.

**Central carbon metabolism.** Out of 115 MAGs, only one Moranbacteria (Bin_419) encode hexokinase, with the potential to phosphorylate glucoses into glucose-6-phosphate. A glycolysis pathway is near complete in most Asgard archaea, Fibrobacteres-Bacteroidetes-

Chlorobi (FBC), Planctomycetes-Verrucomicrobia-Chlamydiae (PVC) group and "others" MAGs (Fig. 3 and Additional file 15: Table S3). Most Parcubacteria and Microgenomates MAGs in the present study lack 6-phosphofructokinase I (*pfk*) and fructose-1,6-bisphosphate phosphatase (*fba*), rendering them an incomplete glycolysis pathway. Bifunctional archaeal fructose-1,6-bisphosphate aldolase (K01622, *FBPA*) was identified in Loki- and Thorarchaeota MAGs, which represents an ancient carbon fixation enzyme in archaea [5]. This enzyme has been identified in Asgard archaea MAGs previously, further supporting Asgard archaea as early evolved microorganisms [6]. Interestingly, Stahlbacteria, Latescibacteria, UBP1, Moranbacteria, Bathyarchaeota and Micrarchaeota MAGs also encode for this enzyme (Additional file 15: Table S3), suggesting these deeply branching lineages retaining primordial metabolisms.

Only 10 MAGs (Heimdallarchaeota, Zixibacteria, GN15, UBP1, Latescibacteria and Uncultured bacterium BMS3Bbin04) harbor a complete TCA cycle, suggesting potential aerobic capacity of these MAGs. Although a complete aerobic kynurenine pathway was identified in Heimdallarchaeota MAGs from brackish-lake sediments in Romania [7], none of the MAGs (including Asgard archaea) in Shark Bay encode a complete kynurenine pathway (Additional file 15: Table S3). This may be due to different environments and abiotic factors shaping different metabolic capacities of resident microorganisms.

Most Archaea, Parcubacteria, and Microgenomates in this study appear to lack genes encoding enzymes (glucose-6-phosphate 1-dehydrogenase, 6-phosphogluconolactonase, 6-phosphogluconate dehydrogenase) involved in the oxidative part of the pentose phosphate pathway (PPP). On the other hand, most of the MAGs encode genes for the non-oxidative part of PPP except that Parcubacteria and Microgenomates lack transaldolase (Fig. 3 and Additional file 15: Table S3). Although Parcubacteria and Microgenomates seem to lack a complete glycolysis and PPP pathway, all MAGs affiliated to these two groups encode glyceraldehyde 3-phosphate dehydrogenase, phosphoglycerate kinase, 2,3-bisphosphoglycerate-independent phosphoglycerate mutase (*gpmI*), enolase (*ENO*), phosphoenolpyruvate synthase (*pps*) and pyruvate kinase. These enzymes facilitate the metabolism of glyceraldehyde 3-phosphate (G3P), the product of the first half of the glycolysis pathway and PPP, to pyruvate. Therefore, it is suggested that these two microbial groups have symbiotic lifestyles requiring hosts with complete PPP pathways or the production of G3P.

**Wood-Ljungdahl Pathway and Methanogenesis**. In agreement with previous studies, genes affiliated with the Wood-Ljundahl pathway were identified in Asgard archaea MAGs [6, 8-11]. All Fibrobacteres and Modulibacteria (KSB3) encode for carbon monoxide dehydrogenase (*cooSF*) and acetyl-CoA synthase (*cdhDE*, *acsB*), allowing these groups to putatively assimilate carbon monoxide (CO) to acetyl-CoA, and are suggested to be carboxydotrophs which are capable of utilising CO [12]. Furthermore, all Asgard archaea (except Thorarchaeota) and Bathyarchaeota MAGs encode for the subunits of acetyl-CoA synthase (*cdhABCDE*), the key enzymes of the Wood-Ljungdahl (WL) pathway. Most of the Asgard archaea MAGs (except Thorarchaeota) encode for a near complete THMPT-WL pathway in which most of the Asgard MAGs lack 5-10-methylenetetrahydromethanopterin reductase (*mer*). One Lokiarchaeota MAG (Bin_186) and Bathyarchaeota (Bin_348) harbor a complete anaerobic $H_2$-dependent THMPT-WL pathway [9] (encoding *fwd, ftr, mtd, mer* and acetyl-CoA synthase). Contrary to the previous studies [6, 8-11], Thorarchaeota does not seem to encode for a THMPT-WL pathway in the Shark Bay systems. On the other hand, only an Aminicenantes MAG (Bin_127) encode for a complete THF pathway but lacking acetyl-CoA synthase (*cdh*), suggesting that this MAG uses tetrahydrofolate (THF) as $C_1$ carrier rather than autotrophic carbon fixation. It is also suggested that Asgard archaea can operate the WL pathway in reverse for organic carbon oxidation [6, 13]. Furthermore, the presence of WL pathways and glycolysis pathways (Fig. 3a), along with acetyl-CoA synthetase (*acs*) and acetate CoA ligase (*acd*) that allows interconversion between acetyl-CoA and acetate, suggests that Asgard archaea are putatively heterotrophic acetogens supporting previous work [11, 14, 15].

Out of all MDM MAGs, only Asgard archaea and Bathyarchaeota MAGs encode for tetrahydromethanopterin S-methyltransferase (*mtr*), which functions to covert methyl-$H_4$MPT to methyl-CoM, with the former as a key intermediate in the HTMPT-WL pathway, and the latter as the key substrate for methanogenesis [16]. However, no methyl-CoM reductase (*mcr*) was identified in any MAGs, therefore it is inconclusive whether MDM MAGs in smooth mats participates in methanogenesis. The lack of methyl-CoM reductase suggests that Asgard archaea are acetogenic rather than methanogenic [9]. This agrees with a previous metagenomics study in Shark Bay [10], in which no *mcr* genes were identified despite analyses indicating high methane production rates [2]. Although a high hydrogenotrophic methanogen population was identified in a 16S rRNA study, and experiments showed that supplying $H_2/CO_2$ resulted in the highest methane production [2], it is still unknown why *mcr* genes were not identified. It may be due to novel genes/mechanisms contributing to methane production in these mats, and it was recently suggested that Cyanobacteria is linked to methane production [106].

**3-hyroxypropionate/4-hydroxybutyrate pathway**. Lokiarchaeota, Thorarchaeota, and Bathyarchaeota encode 2-methylfumaryl-CoA hydratase (*mch*), which suggests putatively their role in the 3-hydroxypropionate cycle. However, this gene may function to assimilate glyoxylate instead of the carbon fixation pathway (Additional file 15: Table S3). Six Lokiarchaeota MAGs harbor both 4-hydroxybutyryl-CoA dehydratase (*abfD*) and enoyl-CoA hydratase, indicating their roles in the carbon fixing 4-hydroxybutyrate (4HB) pathway, which was also found in previous studies [6, 10] (Additional file 15: Table S3). This suggests that Asgard archaea may have an expanded capacity in carbon fixation apart from the Wood-Ljungdahl pathway (WL Pathway).

**CAZy enzymes**. Overall, glycoside hydrolase (GH) genes encoding enzymes that can degrade hemicellulose, animal and other plant polysaccharides are abundant in the FCB group and Asgard archaea MAGs, but are less abundant in other MDM genomes, especially Parcubacteria, Microgenomates, and DPANN archaea (Additional file 6: Figure S5). α-amylases (GH57) were encoded in most of the MAGs, suggesting amylose and starch as one of the most readily available carbon sources in the Shark Bat mats analysed here, which may be one of the main components of the extracellular polymeric substances (EPS) in these mats [10]. This extracellular enzyme allows MDM to degrade starch outside of the cell and subsequent uptake [17]. It is suggested that MDM here have a role in the organic carbon turnover, providing a dynamic carbon source for the microbial mat community [1, 10, 18]. Furthermore, such carbohydrates abundant in extracellular polymeric secretions are highly prevalent in microbial mats, and given EPS degradation is important in fossilization [107], it hints at a potential role of MDM in mat preservation in the fossil record.

Microbial dark matter communities in Shark Bay also harbor CAZys specifically to breakdown celluloses, hemicelluloses, and plant oligosaccharides (Additional file 6: Figure S5). This suggests their ability to digest plant-derived carbohydrates as a carbon source. Furthermore, most of the Parcubacteria encode endoglucanase (GH74), a member in the cellulase family, further suggesting these groups with limited biosynthetic capabilities are able to derive carbon source from plant carbohydrates. Indeed, seasonal cyclones and storms in Shark Bay often bring in large amount of plant biomass from the Faure Sill [19-21], and this may serve to augment carbon sources in the oligotrophic environment of Shark Bay and contribute to the fermentation processes among MDM. Chitinase (GH23) was identified in all MDM groups except Omnitrophica, indicating their ability to degrade chitin, which likely originates from dead eukaryotic cells or molluscs in the area, with the latter frequently found embedded in the microbial mats. The lower range of GH enzymes encoded by Parcubacteria, Microgenomates, Peregrinibacteria, and DPANN archaea suggests these members could scavenge readily degraded carbohydrates through their potential symbiotic hosts or partners.

**Other carbon metabolisms**. Apart from carbohydrate degradation, only Asgard archaea, Bathyarchaeota, and the FCB group bacteria appear to have the genomic capacity to degrade lipids via the beta-oxidation pathway (Fig. 4a, Additional file 4: Figure S3, Additional file 7: Figure S6 and Additional file 15: Table S3), suggesting lipids may not be a common carbon source among microbial dark matter. The ability to oxidise butyryl-CoA to acetyl-CoA allows Asgard archaea to potentially oxidise acetyl-CoA to $CO_2$ through the reverse THMPT-WL pathway, adding to the metabolic versatility of this superphylum [11]. It is suggested that anoxic fermentation of carbohydrates is the main carbon source for the other MDM members in Shark Bay.

Two Moranbacteria (Bin_114, 419) and one Micrarchaeota (Bin_091) MAG encode ATP citrate synthase (*ACLY*), a key gene in the carbon fixing reverse TCA cycle. It was not identified as a major carbon fixation pathway in smooth mats metagenome as described in Wong et al (2018) [10], suggesting MDM may potentially occupy this niche in these mats to maximise energy yield.

Genes encoding dehalogenases are not prominent among MDM MAGs in Shark Bay, indicating that organohalides are likely not a main energy source. Most of the MDM MAGs encode for epoxyqueuosine reductases (*queGH*), but the role of respiring organohalides cannot be determined if they do not encode for reductive dehalogenase domains (IPR028894) [6]. All but two Asgard archaea MAGs (Heimdall-, Thor-, Lokiarchaeota) harbor both epoxyqueuosine reductases and reductive dehalogenase domains, which is in agreement in a previous study [6]. Furthermore, Zixibacteria, KSB1, Bacterium BMS3Bbin04, Aminicenantes (OP8) and Amatimonadetes (OP10) MAGs also encode both epoxyqueuosine reductases and reductive dehalogenase domains (Additional file 15: Table S3). A previously described backbone dataset containing well-established dehalogenases was used to construct a phylogenetic tree to examine if the aforementioned MAGs can respire organohalides [6, 22]. Additional file 20: Table S8 lists the sequences used in the backbone dataset and reductive dehalogenase domain (IPR028894) in this study. Results indicate that Shark Bay MDM MAGs clade with homologous sequences of dehalogenase reductase in Asgard archaea that lack the reductive dehalogenase domains (IPR028894), but not with the *bona fide* reductive dehalogenases identified in previous studies [6, 22, 23] (Additional file 12: Figure S11). Thus it is unclear if the Shark Bay MDM community can respire organohalides, and potentially the different environments between deep subsurface and surface hypersaline microbial mats may have shaped the genomic repertoire of the resident microbial communities.

**Amino acid degradation**. Most of the MDM community in Shark Bay encode for peptidase M28, M50, M20/M25/M40, which are membrane bound peptidases. Furthermore, most MDM MAGs harbor cytoplasmic peptidases family M24, facilitating the putative breakdown of amino acids inside the cell. Metallopeptidase family (M17, M20, M24, M28, M42, M55) and serine peptidases (S9, S33, S58) were identified in most the MDM MAGs (Additional file 15: Table S3), providing the potential for the rare microbiome in Shark Bay not only in scavenging and breaking down oligopeptides, but also polypeptides as a source of carbon, nitrogen, and sulfur.

**RuBisCo**. Almost one third of the MDM genomes encode for ribulose biphosphate carboxylase (RuBisCo) (Fig. 5). Given not all types of ribulose biphosphate carboxylase undergo carbon fixation, a phylogenetic tree was constructed to examine the variety of RuBisCos in these MAGs. The MDM MAGs appear to harbour bacterial and archaeal type III, type IIIa, type IIIb, type IIIc and type IV RuBisCo as described in the main text (Fig. 5 and Additional file 16: Table S4). This suggests that these microorganisms are involved in the AMP nucleotide salvaging pathway, while MAGs harbouring type IV RuBisCo are involved in methionine salvage pathways [24, 25]. Since the RuBisCo in the present study are not classified as type I or type II RuBisCos, MDM MAGs are not involved in photosynthetic carbon fixation. Moreover, none of the RuBisCo-encoding MAGs harbor a complete Calvin-Benson-Bassham cycle (Additional file 15: Table S3). The lack of phosphoribulokinase

(K00855) in the RuBisCo encoding MAGs, an essential enzyme in that converts ribulose 5-phosphate into ribulose 1,5-bisphosphate, also suggests that RuBisCo in Shark Bay MDM are not involved in Calvin-Benson-Bassham cycle. As mentioned in the main text, 22 out of the 32 MAGs with RuBisCo also encode both AMP phosphorylase (*deoA*) and R15P isomerase (*e2b2*) (Additional file 15: Table S3), indicating the potential ability to incorporate $CO_2$ into nucleotide salvaging pathways [26-28]. Ribose-1,5-bisphosphate (R15P) is produced from AMP phosphorylase, subsequently R15P isomerase converts it to ribulose 1,5-bisphosphate (RuBP) [28]. $CO_2$ and $H_2O$ can then be incorporated in RuBP by RuBisCo, resulting in glycerate-3P which then can be fed into the glycolysis [28, 29]. This alternative pathway is suggested to maximise energy yield with MDM that have minimal sized genomes [26].

As mentioned in the main text, one Lokiarchaeota MAG (Bin_186) harbors a type IIIa RuBisCo, which is known to fix $CO_2$ for the synthesis of metabolites using the reductive hexulose-phosphate (RHP) cycle [27]. All the genes necessary for the RHP cycle were identified in this Asgard archaea MAG except phosphoribulokinase (PRK) (Additional file 15: Table S3). PRK is essential for Ribulose-1,6,-biphosphate (RuBP) substrate regeneration, which is critical for the Calvin-Benson cycle. This Lokiarchaeota MAG harbours a complete THMPT-WL pathway (Additional file 15: Table S3), and encodes a fused bifunctional enzyme 3-hexulose-6-phosphate synthase/formaldehyde-activating enzyme (*fae-hps*). These two enzymes together are able to produce methylene-$H_4$MPT from 3-arbino-hexulose-6-phosphate, which is an essential metabolite in the THMPT-WL pathway [27]. Therefore, though the capacity of this Lokiarchaeota for RHP cycling cannot be confirmed as yet, potentially due to an incomplete genome, such an incomplete RHP cycle may serve to replenish $C_1$ carriers in the THMPT-WL pathway. Three Woesearchaeota MAGs (Bin_028, Bin_187, Bin_568) encode for type IIIb RuBisCo, corroborating the findings of a recent study that this type of RuBisCo was only found in DPANN archaea, potentially as a lineage-specific RuBisCo [28]. One interesting finding is that Heimdallarchaeota (Bin_120) contains RuBisCo at the basal position (Fig. 5), suggesting it may possess RuBisCo as an early-evolved form. The wide spread of RuBisCo among MDM in smooth mats suggests ribose, nucleotide-derived sugars and potentially $CO_2$ are fed into the central carbon metabolism to supplement carbon sources, given most of the RuBisCo-harboring MAGs (except the Asgard archaea) encode for an incomplete upper glycolysis pathway and a minimal genomic repertoire.

**Hydrogenases**. $H_2$ was suggested to be an important intermediate in Shark Bay in previous studies. Firstly, a considerable amount of hydrogenotrophic sulfate reducing bacteria were found in smooth mats [1, 2]. Secondly, hydrogenotrophic methanogenesis was found to be the main mode of methane production through rate measurements and a 16S rRNA gene survey [2]. In the current study, 70% (81 out of 115 MAGs) harbor hydrogenases. There are 16 types of hydrogenases divided into 2 groups, which are [NiFe] and [FeFe] respectively. [NiFe] hydrogenases identified in Shark Bay MDM are 1a, 1c, 3b, 3c, 3d, 4a, 4b, 4e, 4g and 4i. [FeFe] hydrogenases identified are *hnd* Group A, A1, Group B, C1, C2 and C3 (Additional file 15: Table S3).

[FeFe] hydrogenases are known to produce $H_2$ and are associated with fermentative $H_2$ production [30-32]. This hydrogenase group was identified in 40 MAGs (Additional file 15: Table S3). Group 3 (3b, 3c, 3d) bidirectional hydrogenases were identified in 62 MAGs, indicating their ability to consume and produce $H_2$. [NiFe] Group 4 (4a, 4b, 4e, 4g, 4i) and [FeFe] Group C (C1, C2, C3) hydrogenases were identified in 16 and four MAGs respectively (Fig. 4). The former has a putative function of ferredoxin-coupled respiration while the latter has a putative function of $H_2$ sensory [33]. However, both roles are unconfirmed and further work is required to characterise their function(s) in the Shark Bay mats.

Parcubacteria and DPANN archaea both encode [NiFe]-3b and [FeFe]-A1 hydrogenases (Additional file 15: Table S3). The co-occurrence of both type of hydrogenases indicate these MAGs potentially undergo fermentative $H_2$-evolution coupled with NADH and ferrodoxin [34]. Other than the suggestion that Woesearchaeota may be in a symbiotic relationship with hydrogenotrophic methanogens, formate can also be used as an electron donor during hydrogenotrophic methanogensis [35-37]. Bacteria affiliated with "others" and Asgard archaea harbor formate dehydrogenase for formate metabolism, though the latter likely channel formate into the Wood-Ljungdahl pathway [8, 11].

Heimdallarchaeota and Thorarchaeota harbor [NiFe] hydrogenase 3b and 3c, which was suggested to work in tandem with WL-pathway, enabling them to grow lithoautotrohpically using $H_2$ as electron donor [6, 9]. Heimdallarchaeota is the only archaeal MAG encoding Group 4b hydrogenase, allowing it to respire formate. It may compensate Heimdallarchaeota to metabolise formate since it is the only Asgard archaea lacking formate dehydrogenase (Additional file 15: Table S3).

**Sulfur and nitrogen cycle**. Genes encoding for a complete dissimilatory sulfate reduction pathway (*dsrAB*, *aprAB*) were identified in Zixibacteria and Zixibacteria order GN15 (formerly classified as a separate phylum: candidate phylum GN15). In addition, genes *dsrEFH* were also identified in Zixibacteria MAGs (except Bin_224 and order GN15). It was reported that *dsrEFH* serve as a role to transfer sulfur to *dsrC*, which in turn is transferred to *dsrAB* acting in the oxidative direction, effectively oxidising sulfite back to sulfate [38-40]. Therefore, this suggests that Zixibacteria in these mats have a role in both dissimilatory sulfur reduction and sulfur oxidation in their hypersaline settings. Other than Zixibacteria, *dsrEFH* were identified in the present study in microbial phyla KSB1, Fibrobacteres, Stahlbacteria (WOR-3), Latescibacteria (WS3), Aminicenantes (OP8), Armatimonadetes (OP10), Coatesbacteria, Eisenbacteria, Poribacteria, Bathyarchaeota and Asgard archaea (Additional file 15: Table S3). These sets of genes were considered restricted to sulfur oxidising bacteria until they were recently identified in Actinobacteria, Candidatus Rokubacteria, and Nitrospirae [41]. This infers an expanded sulfur cycle and the putative roles of sulfur oxidation in the aforementioned MAGs. This is the first report of evidence for Zixibacteria (including GN15, which is formerly classified as Candidate phylum GN15) potentially partaking in dissimilatory sulfate reduction in surface hypersaline settings, and Asgard archaea encoding *dsrEFH*. This expands the lineages taking part in dissimilatory sulfate reduction, which was thought to be carried out exclusively by the following lineages: Deltaproteobacteria, Firmicutes, Thermodesulfobacteria, Actinobacteria, Nitrospirae, Caldiserica and Archaeoglobus [41].

Evidence for nitrogen cycling was examined by searching for key genes in nitrogen fixation, assimilatory and dissimilatory nitrate reduction. Genes encoding nitrogenase (*nifDKH*) were identified in Fibrobacteres (Additional file 4: Figure S3), inferring diazotrophy in this phylum and corroborating findings in a previous study [42]. One Latescibacteria (RBin_199) and Eisenbacteria (Bin_251) encode for a complete dissimilatory nitrate reduction pathway, while nitrite reductase was found in all Lokiarchaeota and Thorarchaeota MAGs (Fig. 4 and Additional file 15: Table S3). The apparent lack of nitrate reductase implies that nitrite does not originate from nitrate reduction. However, the co-occurrence of CO dehydrogenase and nitrite reductase suggests that Asgard archaea may potentially couple CO oxidation to nitrite reduction [43], allowing them to derive energy from an oligotrophic environment (Fig. 4a and Additional file 15: Table S3). Fig. 2 indicates that most MDM in smooth mats do not participate in nitrogen and sulfur cycles, but rather carbohydrate degradation and fermentation.

**Limited metabolic pathways, presence of diversity-generating retroelements (DGRs) and absence of viral defence systems**. Metabolic reconstruction reveals that most of the MDM MAGs have a complete or near-complete glycolysis and pentose phosphate pathways (Fig. 4, Additional file 4: Figure S3, Additional file 7-11: Figure S6-10 and Additional file 15: Table S3). However, the majority of MDM in smooth mats harbour an incomplete tricarboxylic acid (TCA) cycle as mentioned above, indicating the likely preference of an anaerobic lifestyle. Parcubacteria, Microgenomates, Peregrinibacteria, Altiarchaeles, and DPANN archaea all have limited transport and permease proteins for multiple sugars, amino acids, and phosphate (Additional file 15: Table S3). Most of the MAGs associated with MDM were suggested to be living a parasitic or symbiotic lifestyle, especially in anoxic environments [31]. Parcubacteria, Microgenomates, Peregrinibacteria and DPANN archaea in smooth mats do not appear to have specific roles or monolithic metabolic pathways, possessing small genomes which suggests they are early-evolving microorganisms [44].

Such limited metabolic repertoire raises question on how the microbial dark matter community survive under such extreme environment. Based on a previous metagenomics study [10], it is suggested that nutrient cycles are partitioned in these mats in Shark Bay. MDM harbouring scattered genes and incomplete pathways may serve to derive energy by filling in metabolic gaps. For example, as stated above, co-occurrence of CO dehydrogenase and nitrite reductase suggests that Asgard archaea may potentially couple CO oxidation to nitrite reduction [43], allowing them to generate energy for the WL pathway in an oligotrophic environment. Furthermore, Zixibacteria and candidate Zixibacteria order GN15 participate in dissimilatory sulfate reduction, which also potentially participating in sulfur oxidation (except GN15). Overall, MDM in Shark Bay encode multiple genes for carbohydrate degradation and fermentation (Fig. 2, Fig. 3 and Additional file 6: Figure S5). With the majority of the MAGs capable of carbohydrate degradation and fermentation (Fig. 3 and Additional file 6: Figure S5), it is proposed that these microorganisms may have important roles in carbon cycling, such as recycling dead cells and microbial biomass, or even degraded plants [45-48]. As mentioned above, seasonal cyclones and storms in Shark Bay often bring in large amount of plant biomass from the Faure Sill [19-21], and this may serve to augment carbon sources in the oligotrophic environment of Shark Bay and contribute to the fermentation processes among MDM.

Given the minimal metabolic capacities and a proposed symbiotic lifestyle of the MDM in the Shark Bay mats [14, 44], analyses of diversity-generating retroelements (DGR) in the Shark Bay MAGs was undertaken. DGRs enable microbes to modify DNA sequences and proteins, which usually targets proteins involved in surface attachment and defence [14, 49, 50]. By employing the mechanism of mutagenic homing, DGRs are capable to mutate surface proteins with an infinite range of protein variants, acting as an agent for cell-cell attachment and dynamic host responses [50, 51]. This facilitates host-dependent microorganisms to attach to their hosts' surfaces for a symbiotic lifestyle. Most of the DGRs were identified in Parcubacteria and DPANN archaea, which may link to the minimal metabolic capacities they harbor as illustrated in the current and previous studies [14, 50]. However, in the present study, DGRs were also identified in Asgard archaea (Lokiarchaeota; RBin_035, RBin_125, Bin_186), which has not been reported before. Despite having versatile metabolisms (WL pathway, fatty acid/amino acid degradation, nucleotide salvaging pathways, putative lithoautotrophy, heterotrophic acetogenesis and light sensing rhodopsin), this may indicate Asgard archaea once resided in energy-limited environments [49].

Virus defence systems CRISPR, BREX and DISARM were identified in MAGs affiliated mainly to Asgard archaea, FCB, and PVC groups (Fig.1 and Fig. 2). Only one Lokiarchaeota MAG (RBin_125) encode a full set of *dndCDEA-pbeABCD* as a novel type of DNA phosphorothioation-based viral defence system (Additional file 15: Table S3) [52]. Parcubacteria MAGs are almost devoid of any viral defence systems except for a Portnoybacteria MAG (Bin_561), despite a recent study describing an abundant viral community associated with the Shark Bay mats suggests the potential for viral predation [53].As mentioned in the main text, the absence of any identified virus defence systems may be due to MDM acting as 'viral decoys', avoiding autoimmunity and to avoid high energetic cost to maintain such systems, as they harbor limited metabolic capacities [54-56]. On the other hand, frequent viral infections may influence genome dynamics due to an evolutionary arms-race between viruses and hosts and could thereby contribute to increased rates of evolution of microbial dark matter [57, 58], or even gain of function. Such recombination events through HGT were found to contribute to the formation of genomic islands that are linked by common functional and evolutionary themes [59]. Examples include virulence islands, polymorphic toxins, defence islands, and integrated elements [60-65]. This may be a putative mechanism as to how MDM acquire genes required to survive in certain extreme environments despite possessing minimal size genomes. As described in the main text, it is suggested that synergy between presence of DGRs and absence of viral defence systems results in rapid screening and acquisition of biological functions for survival.

**Early-evolved genes and ancient traits in Shark Bay mats**. Heimdallarchaeota (Bin_120) encodes for a RuBisCo at a basal position in the phylogenetic tree (Fig. 5), suggesting an early-evolved form of RuBisCo in Shark Bay, and supporting the evidence that Heimdallarchaeota as an early branched lineage [66, 67]. Asgard archaea in the Shark Bay mats can potentially encode for both THF- and THMPT-WL pathways (Fig. 3a and Additional file 15: Table S3). Both THMPT- and THF-WL pathways were also identified in Lokiarchaeota in other ecosystems [8, 9, 11, 67], suggesting a versatile metabolism in Asgard archaea. The CODH/ACS complex involved in WL pathways are hypothesised as an early-evolved complex, further supporting that Asgard archaea as an early-branching lineage [68]. Moreover, THMPT-WL pathways coupled with hydrogenotrophic methanogenesis (HG) were thought to be a trait in the last universal common ancestor of archaea [11, 16, 37], with HG found as the main methane production mode in Shark Bay [2]. Additional evidence will be needed to trace the evolutionary history of WL pathways and how they converge in Asgard archaea. However, it is suggestive of ancient traits present in the modern Shark Bay systems.

MDM MAGs also encode for arsenic resistance despite their minimal genomes. It is suggested that arsenic resistance genes are ancient artefacts, as microorganisms present in the

Precambrian Earth were believed to couple arsenic metabolism with carbon and nitrogen cycles [69, 70]. This further suggests the potential for aspects of Shark Bay mat genomes to provide insights into life on the Precambrian Earth.

The identification of DGRs in reduce-sized genomes suggests that protein evolution was accelerated to facilitate adaptation to selective pressures and symbiotic associations [50]. Some of these retroelements are suggested to have evolved useful functions to benefit their hosts and integrated into the bacterial and archaeal hosts [71]. It was proposed that certain early-evolved biological functions encoded by the DGRs were retained in the hosts therefore further studies on the DGRs in Parcubacteria and DPANN archaea could potentially act as a window to the past [71].

**Isoprenoid biosynthesis pathway and lipid divide in microbial dark matter**.

As Parcubacteria and DPANN archaea were suggested to be early evolving microorganisms [14], isoprenoid lipid biosynthesis pathways were examined in the present study to investigate the 'lipid divide' of bacteria and archaea [44]. Bacteria usually undergo the methylerythritol phosphate (MEP) pathway [108], while the mevalonate (MVA) pathway is predominantly found in archaea and eukaryotes, and has only been found in a few bacteria [109, 110]. A near-complete bacterial MEP pathway was identified in a Woesearchaeota MAG (Bin_434), which was only recently found in another Woesearchaeota residing in deep subsurface environments [44].

Apart from a near-complete bacterial MEP pathway being identified in a Woesearchaeota, isopentenyl phosphate kinase (*ipk*), a gene affiliated to the archaeal MVA pathway was identified in two KSB1 and two Pacebacteria MAGs in the present study (Additional file 15: Table S3). A complete eukaryotic MVA pathway (with phosphomevalonate kinase [*PMK*], diphosphomevalonate carbocylase [*MVD*], isopentenyl diphosphate isomerase [*IDI*]), was found in a Nealsonbacteria (Bin_162) and Woesearchaeota (Bin_274) MAG, and near complete eukaryotic MVA pathways were also found in Lokiarchaeota, Dojkabacteria (WS6), Dependentiae (TM6), and FCB group MAGs (Additional file 15: Table S3). Genes encoding eukaryotic MVA enzyme *IDI1* was also identified in DPANN MAGs, which was also reported in a recent survey [44]. The eukaryotic MVA pathway identified in both bacteria and archaea was suggested to arise not as a result of horizontal gene transfer, but rather as a trait of the last common ancestor (cenancestor) of bacteria and archaea [72]. Findings in the present study reinforces the suggestion that the eukaryotic MVA pathway is a trait of the last common ancestor (cenancestor) of bacteria and archaea [44, 72]. However, the discovery of the MEP pathway in Woesearchaeota suggests the possibility of horizontal gene transfer. The reported distribution of MVA and MEP pathways blurs the distinct "lipid divide" and changed the prior concept that the MEP pathway can only be found in bacteria [44].

**Eukaryotic signature proteins (ESPs)**.

The emergence of the eukaryotic cell is one of the most controversial issues in evolutionary biology. The presence of eukaryotic signature proteins (Additional file 3: Figure S2), proteins in eukaryotes with no significant homologues in archaea or bacteria, has led some to argue that eukaryotes emerged from complex cells distinct to bacteria, archaea or modern-day eukaryotes, terming these cells chronocytes [73]. However, an abundance of ESP has been recently reported in the superphylum of Asgard archaea [67, 76], suggesting that Asgard archaea possess complex eukaryotic-like characteristics and hinting at a close evolutionary relationship between Asgard archaea and eukaryotes.

To assess the evolutionary relationship between eukaryotes and the Asgard archaea of Shark Bay, the MAGs were screened for ESP [7, 67, 73-76] by annotating against the PFAM/TIGRFAM databases using Interproscan5 [77] and the KEGG database using GhostKoala [78], with protein homology confirmed using HHpred [79] and BLAST [80]. In keeping with previous studies [67, 76], the MAGs of Asgard archaea were found to encode ESP, including those involved in cytoskeleton dynamics, information processing, trafficking machinery, signalling systems and N-linked glycosylation (Additional file 3: Figure S2). The MAGs of Shark Bay Asgard archaea were found to encode an abundance of actin family proteins, as previously described [67, 76]. In terms of information processing genes, five new ESPs were identified in the MAGs of Shark Bay Asgard archaea. Amongst these new ESPs was the eukaryotic elongation factor 1-β (Bin_186, Bin_204, Bin_229, Bin_485, RBin_125, Bin_478, RBin_111, Bin_120), the proteasome regulatory particle subunit 11 (Bin_186, Bin_204, Bin_229, Bin_342, RBin_035, RBin_125), subunit 5 of the COP9 signalosome complex (Bin_204, RBin_035), subunit 2 of the transcription initiation factor TFIIH (Bin_229) and a 18S rRNA methyltransferase (Bin_186, Bin_204, Bin_229, Bin_342, Bin_485, RBin_035, RBin_125). In line with previous work [67, 76], the catalytic (Alg13) subunit of the N-linked glycosylation protein UDP-GlyNAc transferase was identified (Bin_478, RBin_111), indicating Asgard archaea possess eukaryotic-like protein modification systems. The Shark Bay Asgard archaea were also found to be enriched for eukaryotic-like signalling systems, including GTP binding proteins, similar to what has been reported for other Asgard archaea [67, 76]. Functional classification of these GTP binding proteins against the KEGG database found that these GTP binding proteins belong to the

ARF (all Asgard MAGs), RAB (all Asgard MAGs), RAN (Bin_204, RBin_035) and RAS (all Asgard MAGs) families, whereas only the ARF and RAS families had been previously described in Asgard archaea [67, 76]. Calmodulin (Bin_485), a eukaryotic dual specificity protein tyrosine phosphatase (Bin_485, RBin_125) and protein phosphatase 1 regulatory subunit 7 (Bin_204, Bin_229, Bin_342, Bin_485, RBin_035, RBin_125) were also identified in the MAGs of Asgard archaea for the first time, suggesting the possession of eukaryotic-like signalling systems.

**Environmental adaptation**. Evidence for salinity adaptation was first examined by delineating genes involved in synthesis and importation of glycine betaine, trehalose and ectoine, as these mechanisms were shown to be the preferred mode for osmoadaptation in hypersaline environment (68 PSU) of Shark Bay [10, 81, 82]. Osmoprotectant permease proteins and glycine betaine transporters are almost exclusively identified only in FCB group MAGs (Fig. 2 and Additional file 4: Figure S3). Moreover, besides the FCB group, only two Elusimicrobia MAGs encode for the complete trehalose biosynthesis pathway (Additional file 10: Figure S9 and Additional file 15: Table S3). Hence, compatible solute accumulation as an osmoadaptative strategy does not appear to be common among MDM in smooth mats. However, potassium uptake proteins and $Na^+$ symporters were found in smooth mat MDM MAGs except for Microgenomates, Parcubacteria and an uncultured archaea (Fig. 4, Additional file 4: Figure S3, Additional file 7-11: Figure S6-10 and Additional file 15: Table S3), indicating the rare biosphere likely adapt a “salt in” strategy, retaining osmotic balance by maintaining high intracellular salt concentrations [83, 84].

Out of the 115 microbial dark matter MAGs, 88 encode for copper resistance genes and over 60% of these MAGs harbour arsenic resistance genes (Figs. 2-4, Additional file 4: Figure S3, Additional file 7: Figure S6 and Additional file 8-11: Figure S7-S10) , suggesting that despite having minimal sized genomes, MDM appear to have adapted to the high copper concentrations in Shark Bay as described in a previous study [10]. Phosphorus intake genes were investigated given the extremely low phosphorus concentration measured in Shark Bay as stated in Wong et al (2018) [10] and previous studies [85-87]. However, phosphorus intake genes (*pho*, *phn* and *pst*) were not detected in Parcubacteria, Microgenomates, and any DPANN archaea MAGs. Furthermore, polyphosphonate associated genes were not identified in Parcubacteria, Microgenomates, and all archaeal MAGs (Additional file 15: Table S3). It was suggested that archaea could utilise their own DNA or extracellular DNA (eDNA) as a phosphorus source [88], and the RuBisCo-bearing MAGs may potentially scavenge free phosphate groups from nucleotides upon the AMP pathway [26, 28]. MDM acting as 'viral decoys' for their host can also putatively scavenge phosphorus from degraded viral DNA.

Genes encoding Type IV pili was found in all groups of bacterial MDM (Fig. 4, Additional file 4: Figure S3, Additional file 7: Figure S6 and Additional file 8-11: Figure S7-S10 and Additional file 15: Table S3). This indicates that microorganisms associated with MDM have the potential ability for processes such as adhesion, motility, protein secretion, and DNA uptake [89]. Archaeal type IV pili (archaellum) are known to be present in a range of archaea [90]. Interestingly, the DUF2341 domain that is associated with archaeal type IV pilli was also found in all Fibrobacteres MAGs in the present study, which is possibly due to horizontal gene transfer. Archaeallum ATPase (*flaI-A*) and membrane platform protein (*flaJ-A*) were found in most archaeal MAGs, however the other archaellum components such as *flaC/E/D* are absent (Additional file 15: Table S3). This may also explain the widespread abundance of RuBisCo, type IV pili, and archaellum in MDM, as they facilitate DNA uptake potentially from eDNA or viral DNA as an extra carbon and phosphorus source [91, 92]. Type IV pili and archaellum also allows interactions with neighbouring microorganisms for communication though no AHL synthases were found, indicating either the absence of quorum sensing by the *lux* mechanism [14, 93], or alternative communication molecules are employed. It is suggested that the archaellum work in concert with DGRs for surface attachments of their hosts, and compensate for the apparent lack of transporters in CPR bacteria and DPANN archaea [50]. Given the diverse nature of the archaellum, they may also have a role in biofilm formation in the mats [93, 94]. Type IV pili and archaellum can also give microorganisms motility, which may facilitate movement between hosts, energy sources, or even niches.

**A conceptual ecological model of MDM in Shark Bay microbial mats**. Although microbial dark matter MAGs appear to have minimal genome size and limited metabolic capabilities, they have been found in various oligotrophic environments such as hydrothermal sediments [95, 96], terrestrial subsurface aquifers [46, 93, 97, 98], deep sea “dead zone” [99] and hypersaline microbial mats as in this study. Apart from the adaptation strategies discussed above, it is proposed that MDM’s main lifestyle is parasitic or symbiotic with other microbial hosts as suggested previously [14, 97, 100].

Previous studies have shown that archaeal MDM (especially Asgard and DPANN archaea) contain genomic contents with very low detectable similarity to the current databases [59]. These sequences (from 30% to 80%) are labelled as the ‘twilight zone’ of sequence similarity to hypothetical proteins with unknown functions [59, 101]. This genomic dark matter may encode for genes that contributes to the survival and metabolic capacity in extreme environments such as Shark Bay. It is proposed that the Shark Bay mats harbour some microorganisms and functional genes that may be relics from early Earth. In addition, it should be noted that within microbial dark matter clades, there is an abundance of genes that are un-annotatable with current databases, and indeed up to 50% of the genes in the Shark Bay MDM were unannotated. These unknowns represent a wealth of data on the MDM in modern mats that could be used for further analysis and give added insights into the roles of these enigmatic groups, thus 'illuminating' microbial dark matter [102]. This may also indicate smooth mat archaea and deep branching lineages retain primordial metabolism that utilise $H_2$, $CO/CO_2$ as biosynthetic starting material [5].

Taken together, microbial dark matter in Shark Bay are proposed to have an ecological role in anoxic carbon and hydrogen transformation. Building on the existing ecological model in Shark Bay [10], apart from Deltaproteobacteria, Chloroflexi and Gemmatimonadetes taking part in dissimilatory sulfate reduction, Zixibacteria and Zixibacterial order GN15 are also proposed to be involved in this pathway (Fig. 6). Partitioning of nitrogen and sulfur cycles were suggested in a previous study, in which these cycles maybe coupled with CO oxidation [43]. To adapt to the hypersaline environment, MDM in Shark Bay adapts the 'salt-in' strategy instead of the prominent glycine betaine accumulation strategy found previously [10, 81, 82]. Photo-degradation may occur, resulting in CO production from organic carbon, which is oxidisied as an alternative carbon source for energy conservation [10, 84, 103]. The resulting $CO_2$ can be potentially assimilated through the AMP nucleotide salvaging pathway, with ribose substituting hexose at the upper part of glycolysis, maximising energy yield. The extensive hydrogenases identified suggests high turnover rate of hydrogen, potentially forming consortium with hydrogenotrophic methanogens by providing $H_2$ in exchange of nutrients [104]. Ribose, $CO_2$/CO and $H_2$ are suggested to be prominent currencies among Shark Bay mat novel uncultured microbiomes.

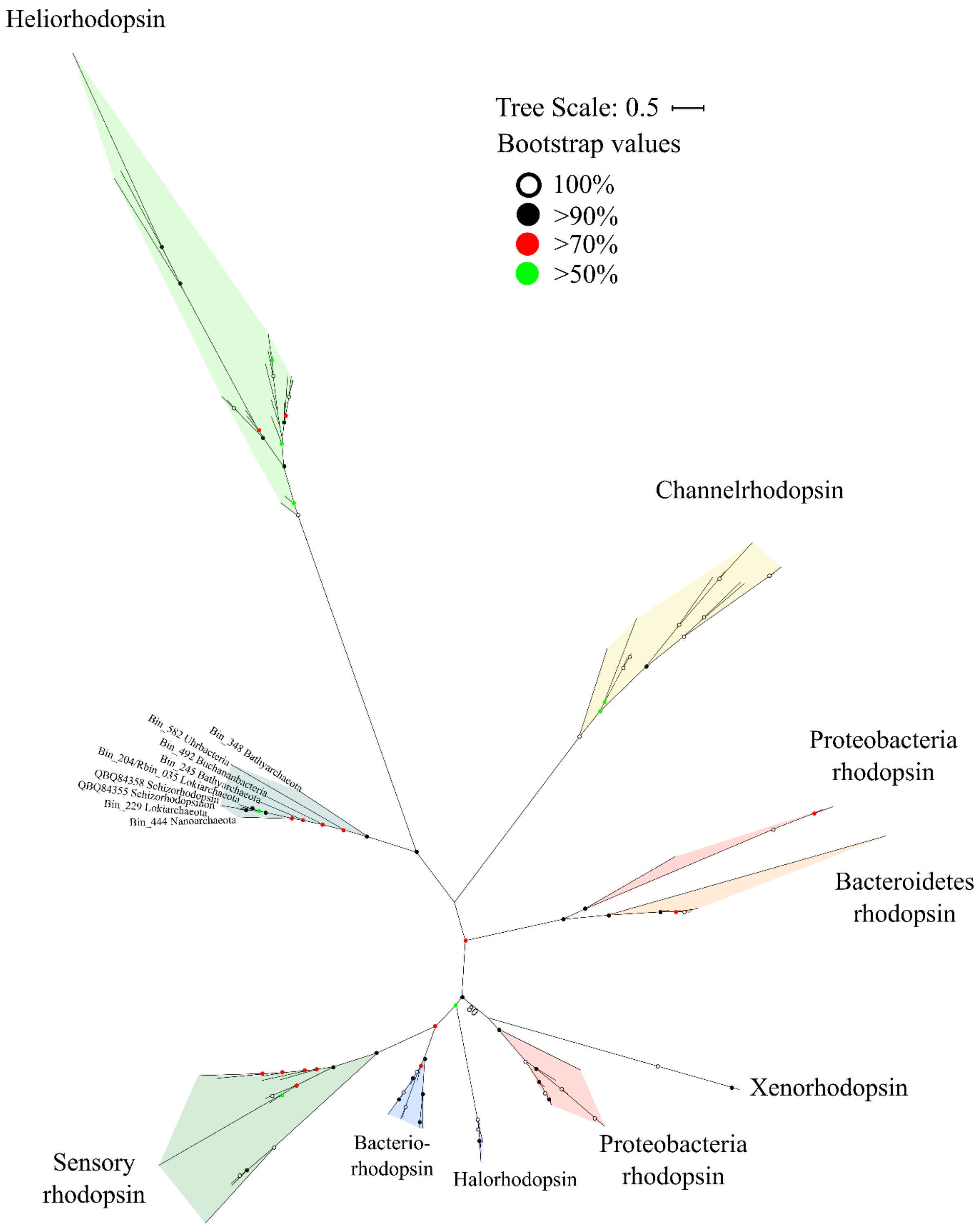


**Additional file 2: Figure S1. Unrooted maximum-likelihood phylogenetic tree of putative rhodopsin in Shark Bay MDM MAGs.** Maximum-likelihood phylogenetic tree constructed with rhodopsin gene found in the MDM MAGs with 1000 bootstrap replications. Lokiarchaeota, Bathyarchaeota, Uhrbacteria, Buchananbacteria and an unclassified archaeon encode rhodopsin clustered in the same group with the novel, recently discovered schizorhodopsin (7). Circular dots of different colors represent bootstrap values. Rhodopsin sequences in this study, reference sequences and BLAST results are listed in Additional file 15: Table S3.

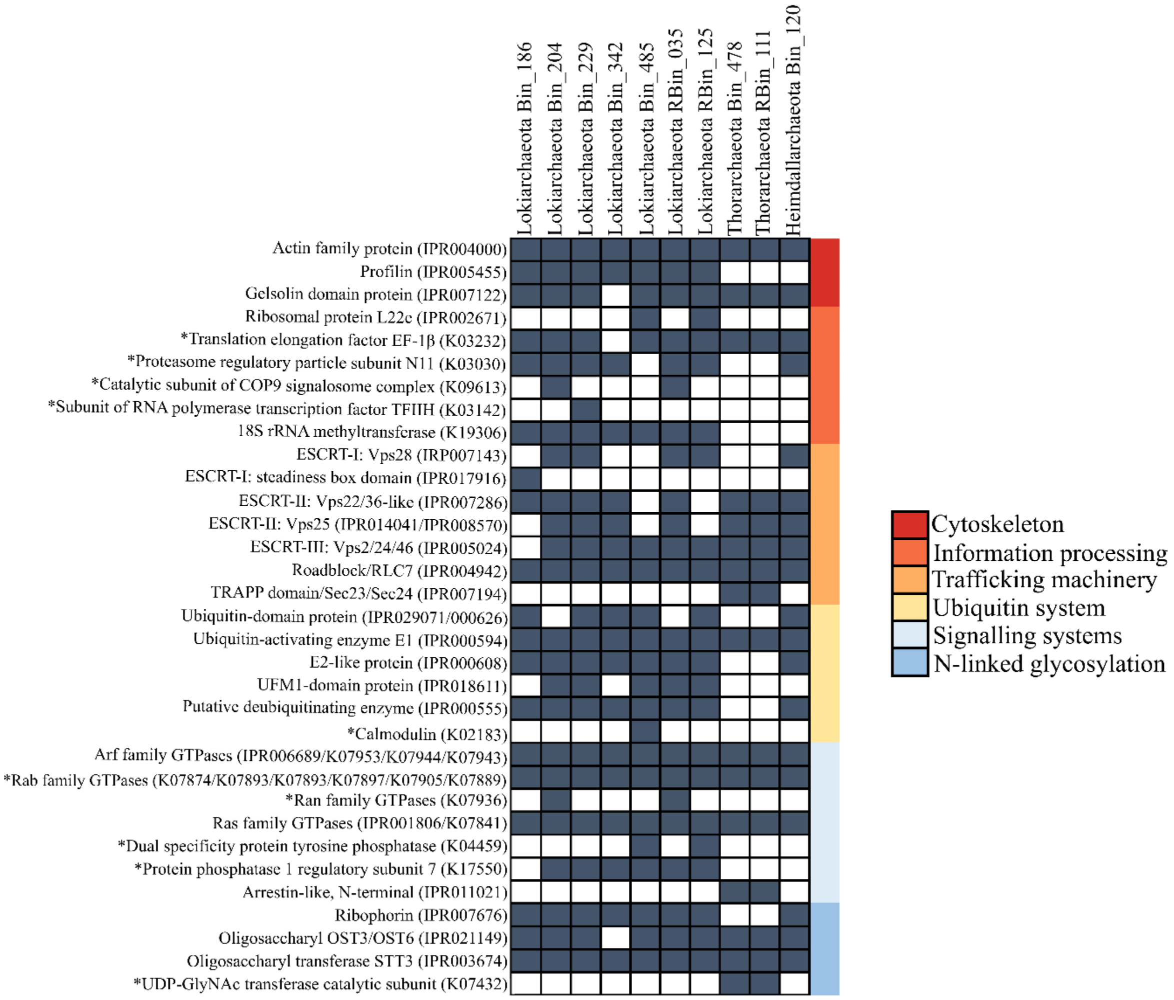


**Additional file 3: Figure S2. Eukaryotic Signature Proteins (ESPs) in the MAGs of Asgard archaea.** MAGs were annotated using InterProScan [77] and GhostKoala [78] and confirmed using HHpred [79] and BLAST [80]. Shark Bay Asgard archaea were found to contain ESP likely involved in cytoskeleton dynamics, information processing, trafficking machinery, signalling systems as well as eukaryotic-like N-linked glycosylation. * indicates newly identified ESP.

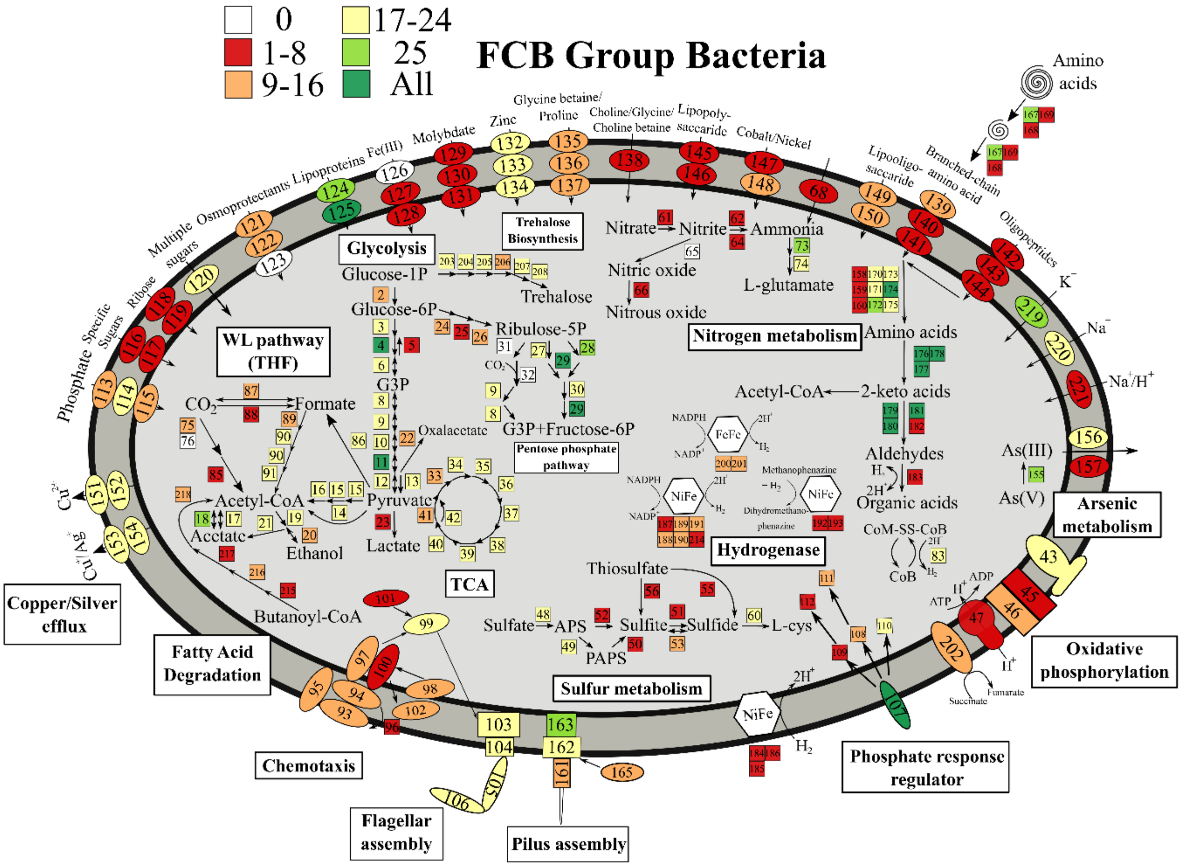


**Additional file 4: Figure S3. Metabolic potential of FCB (Fibrobacteres-Chlorobi-Bacteroidetes) group bacteria.** A metabolic map summarising the genomic potential and metabolic capacities of the 26 MAGs affiliated with the FCB group. Numbers represent specific genes in given pathways and the corresponding genes are listed in Additional file 15: Table S3. Different colors in the square boxes represent different numbers of MAGs encoding the genes, while white square boxes indicate the absence of the genes. TCA, tricarboxylic acid cycle; THF, tetrahydrofolate; WL pathway, Wood-Ljungdahl pathway; PAPS, 3’-phosphoadenylyl sulfate; APS, Adenylyl sulfate.

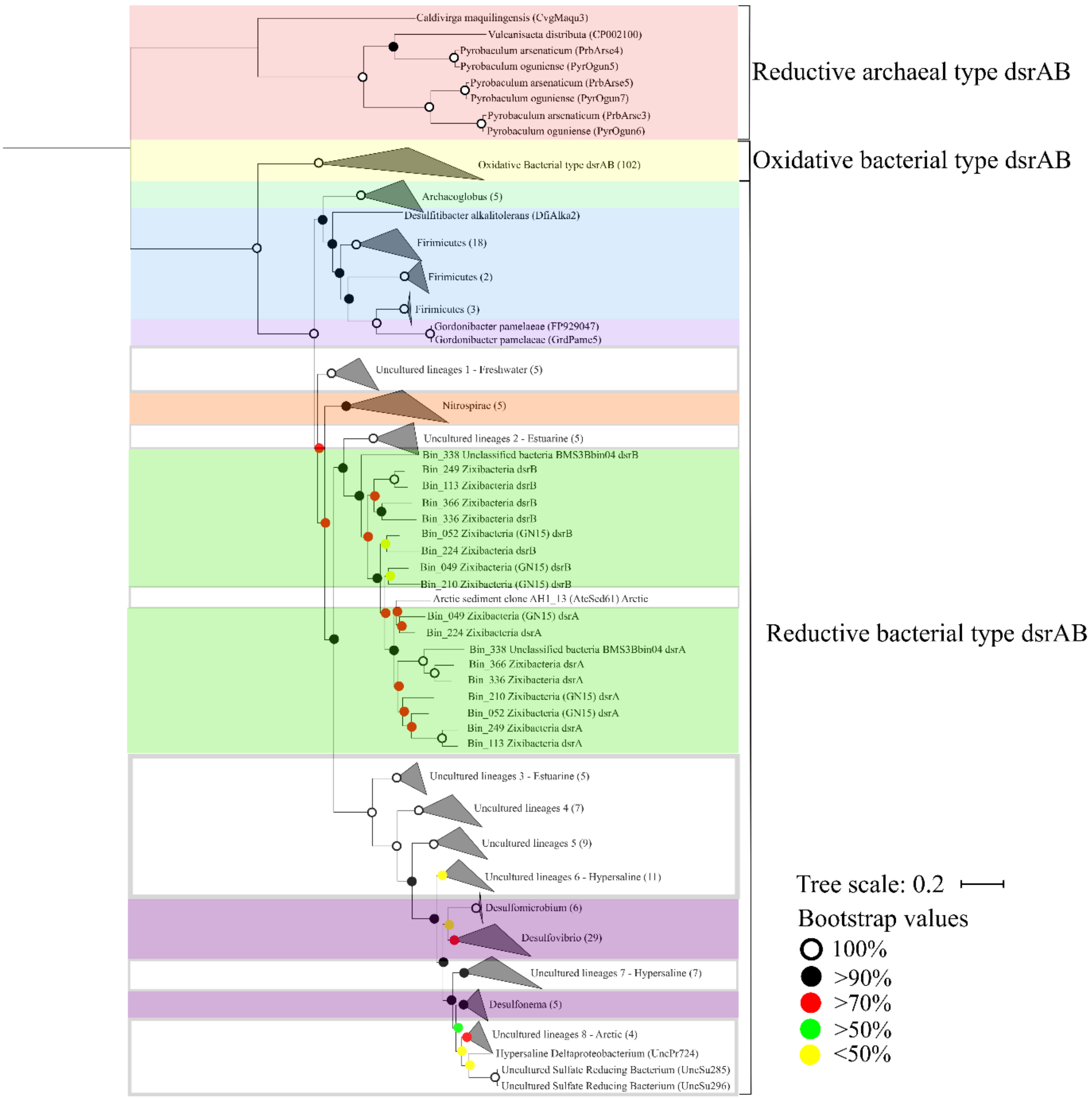


**Additional file 5: Figure S4. Maximum-likelihood phylogenetic tree of *dsrAB* in Shark Bay MDM MAGs.** Maximum-likelihood phylogenetic tree was constructed with reference *dsrAB* sequences from the dsrAB database [105], with 1000 bootstrap replications. *dsrAB* genes found in the present study are classified as reductive bacterial type *dsrAB* and are highlighted in green. Circular dots of different colors represent bootstrap values. *dsrAB*

sequences found in the MDM MAGs are listed in Additional file 19: Table S7. Branches shaded red indicates reductive archaeal type *dsrAB*, yellow shade indicates oxidative bacterial type *dsrAB*, light green indicates Archaeoglobus lineages, light blue indicates Firmicutes lineages, light purple indicates Actinobacteria lineages, orange represents Nitrospirae lineages, purple represents Deltaproteobacteria lineages, green represent *dsrAB* in the present study and no shades represent uncultured/environmental lineages.

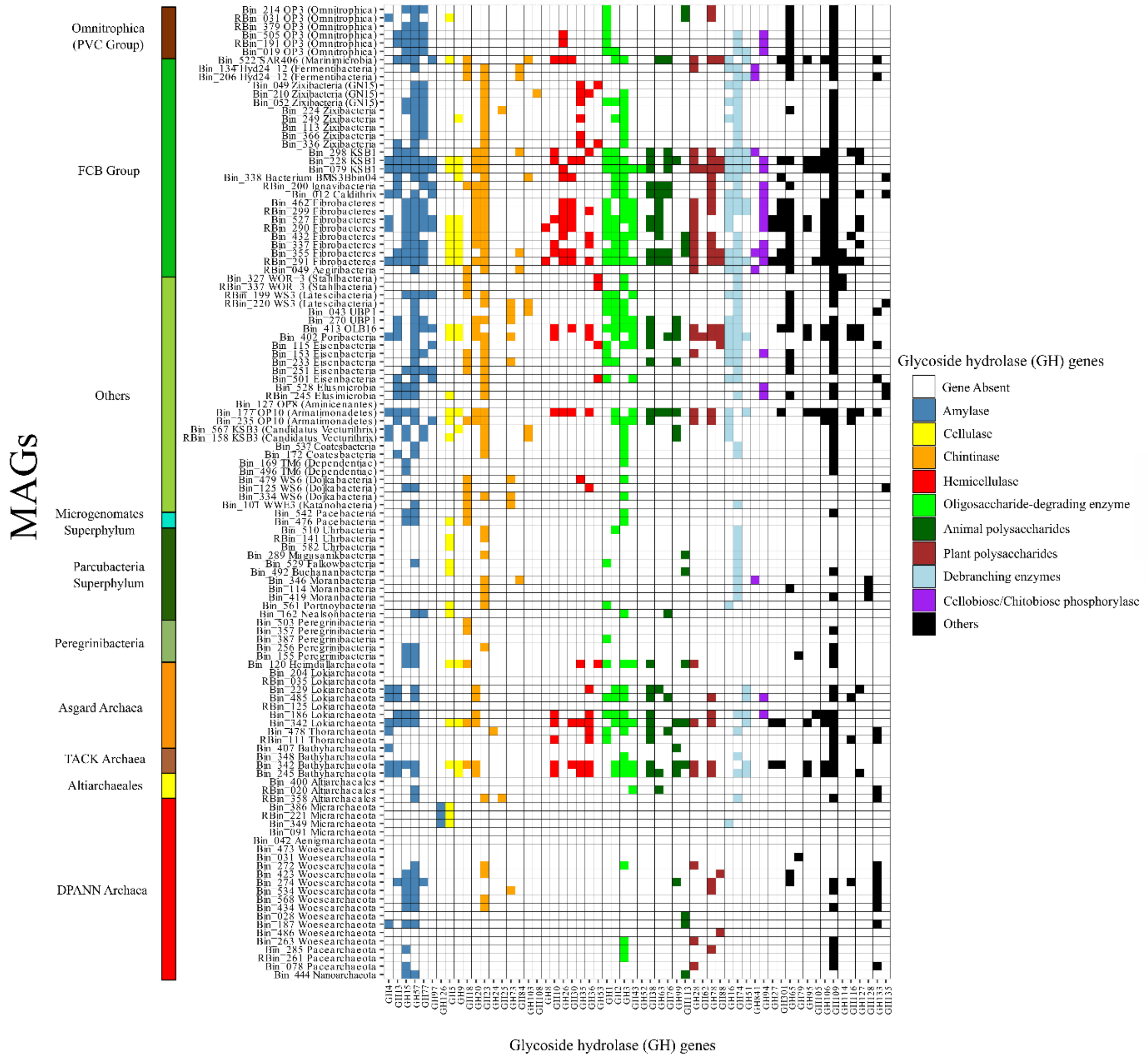


**Additional file 6: Figure S5. Color-coded table indicating major carbohydrate-active enzymes (CAZy) in MDM MAGs.** X-axis indicates different types of glycoside hydrolase (GH) genes in the CAZy database and y-axis represent MAGs of microbial dark matter.

White indicates absence of GH genes in the MAGs. Color panel on the left represents different groups of MDM MAGs according to Fig. 1.

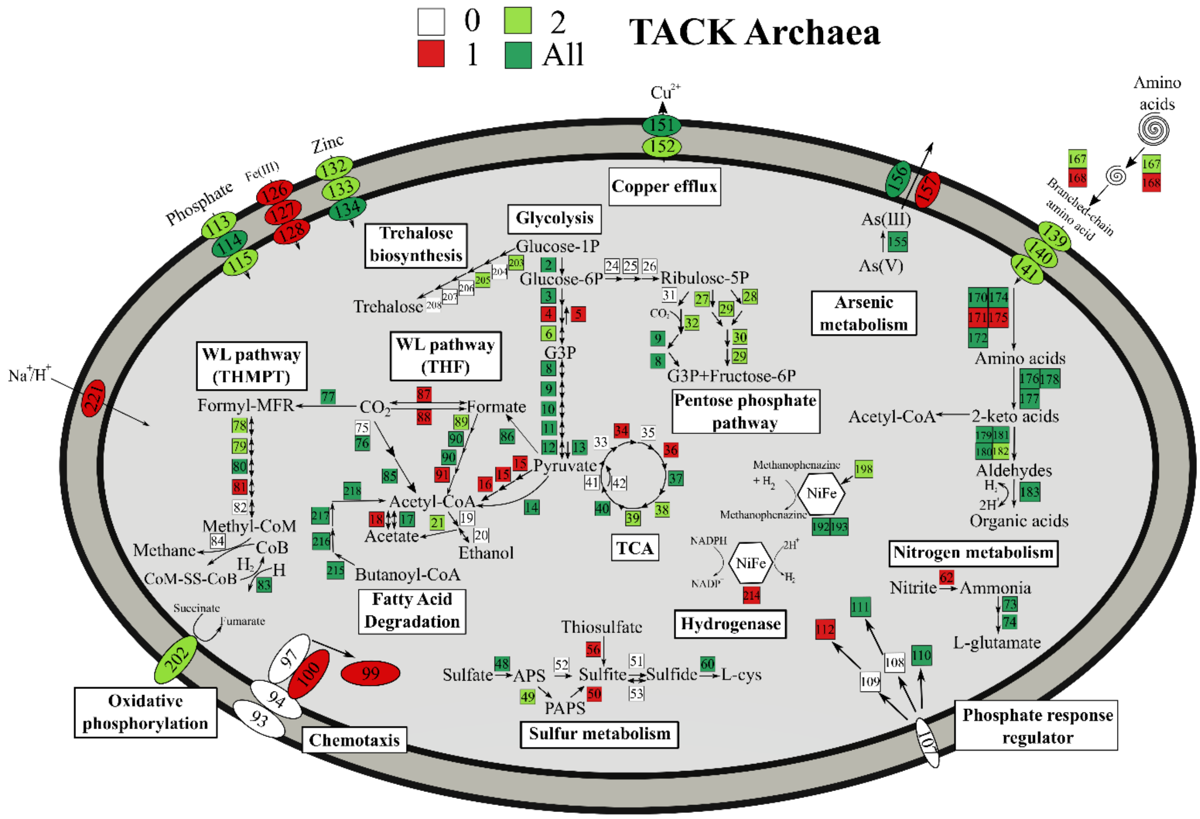


**Additional file 7: Figure S6. Metabolic potential of Bathyarchaeota (TACK archaea).** A metabolic map summarising the genomic potential and metabolic capacities of the 3 MAGs affiliated with TACK archaea. Numbers represent specific genes in given pathways and the corresponding genes are listed in Additional file 15: Table S3. Different colors in the square boxes represent different numbers of MAGs encoding the genes, while white square boxes indicate the absence of the genes. TCA, tricarboxylic acid cycle; THF, tetrahydrofolate; THMPT, tetrahydromethanopterin; WL pathway, Wood-Ljungdahl pathway; PAPS, 3’-phosphoadenylyl sulfate; APS, Adenylyl sulfate.

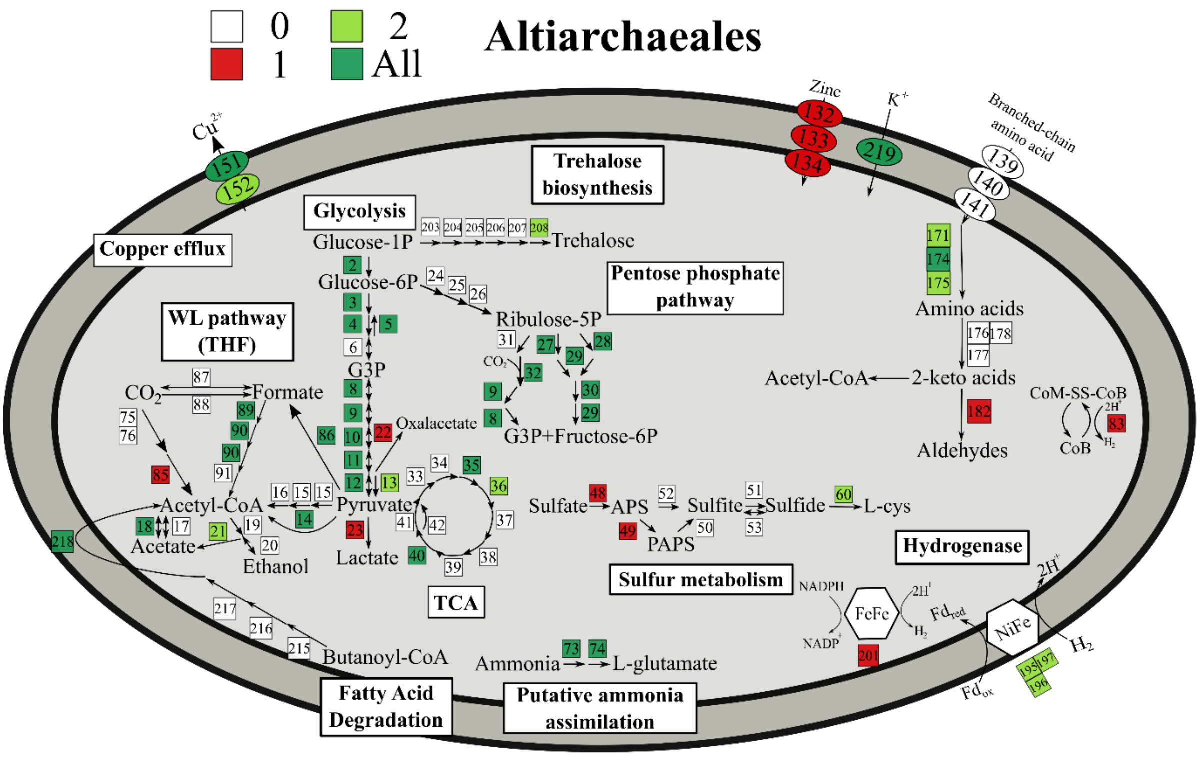


**Additional file 8: Figure S7. Metabolic potential of Altiarchaeales.** A metabolic map summarising the genomic potential and metabolic capacities of the three MAGs affiliated with Altiarchaeales. Numbers represent specific genes in given pathways and the corresponding genes are listed in Additional file 15: Table S3. Different colors in the square boxes represent different numbers of MAGs encoding the genes, while white square boxes indicate the absence of the genes. TCA, tricarboxylic acid cycle; THF, tetrahydrofolate; WL pathway, Wood-Ljungdahl pathway; PAPS, 3’-phosphoadenylyl sulfate; APS, Adenylyl sulfate.

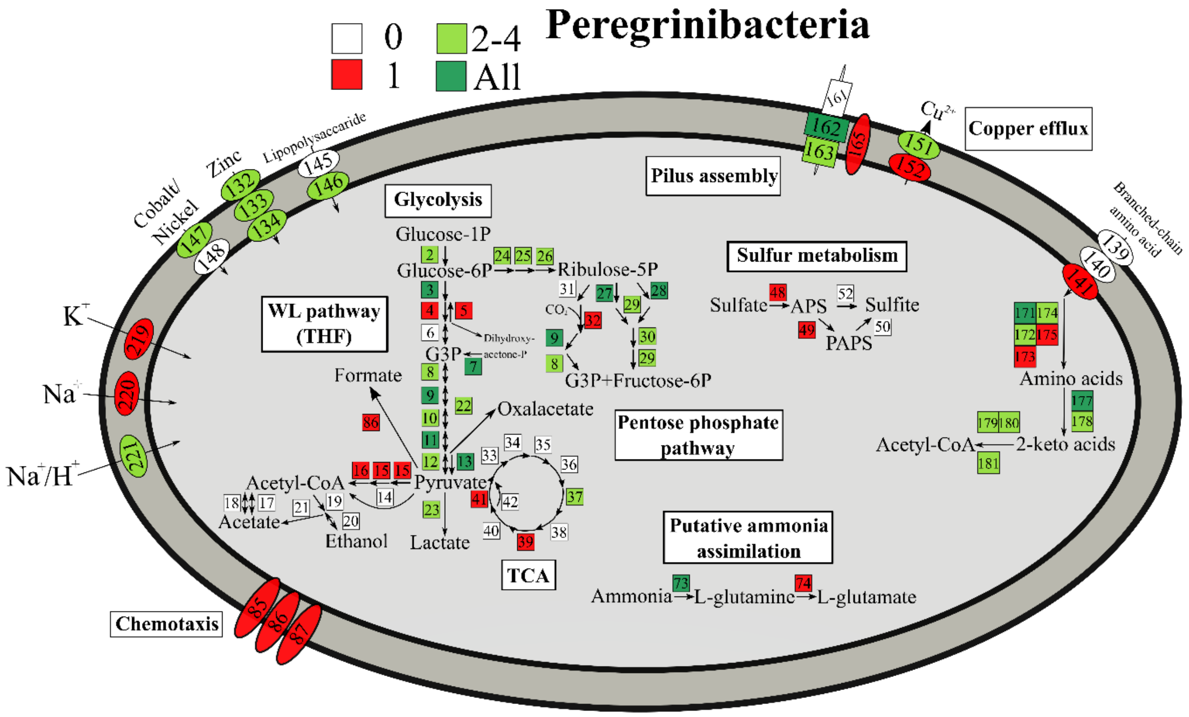


**Additional file 9: Figure S8. Metabolic potential of Peregrinibacteria.** A metabolic map summarising the genomic potential and metabolic capacities of the 5 MAGs affiliated with Peregrinibacteria. Numbers represent specific genes in given pathways and the corresponding genes are listed in Additional file 15: Table S3. Different colors in the square boxes represent different numbers of MAGs encoding the genes, while white square boxes indicate the absence of the genes. TCA, tricarboxylic acid cycle; THF, tetrahydrofolate; WL pathway, Wood-Ljungdahl pathway; PAPS, 3’-phosphoadenylyl sulfate; APS, Adenylyl sulfate.

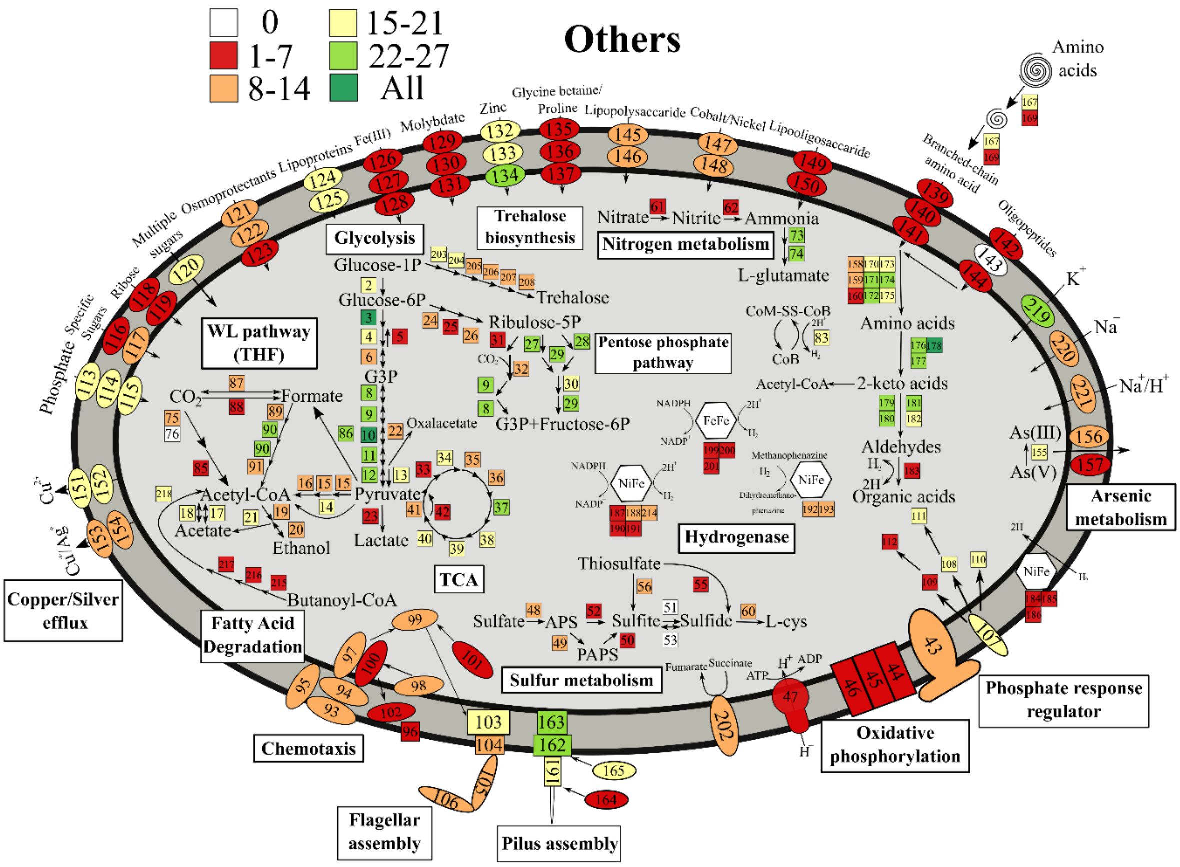


**Additional file 10: Figure S9. Metabolic potential of other MDM bacteria.** A metabolic map summarising the genomic potential and metabolic capacities of the 28 MAGs affiliated with other MDM bacteria. Numbers represent specific genes in given pathways and the corresponding genes are listed in Additional file 15: Table S3. Different colors in the square boxes represent different numbers of MAGs encoding the genes, while white square boxes indicate the absence of the genes. TCA, tricarboxylic acid cycle; THF, tetrahydrofolate; WL pathway, Wood-Ljungdahl pathway; PAPS, 3'-phosphoadenylyl sulfate; APS, Adenylyl sulfate.

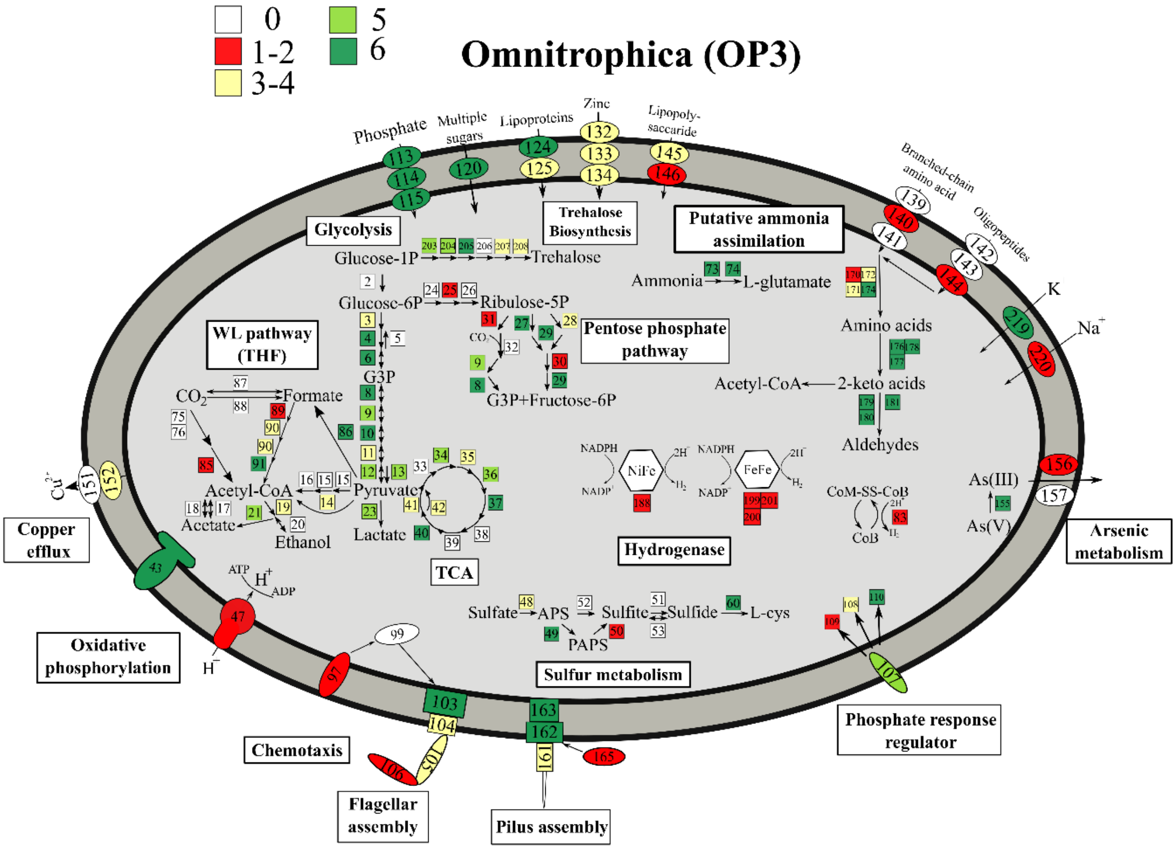


**Additional file 11: Figure S10. Metabolic potential of the PVC (Planctomycetes-Verrucomicrobia-Chlamydiae) group bacteria.** A metabolic map summarising the genomic potential and metabolic capacities of the six MAGs affiliated with Omnitrophica (OP3). Numbers represent specific genes in given pathways and the corresponding genes are listed in Additional file 15: Table S3. Different colors in the square boxes represent different numbers of MAGs encoding the genes, while white square boxes indicate the absence of the genes. TCA, tricarboxylic acid cycle; THF, tetrahydrofolate; WL pathway, Wood-Ljungdahl pathway; PAPS, 3’-phosphoadenylyl sulfate; APS, Adenylyl sulfate.

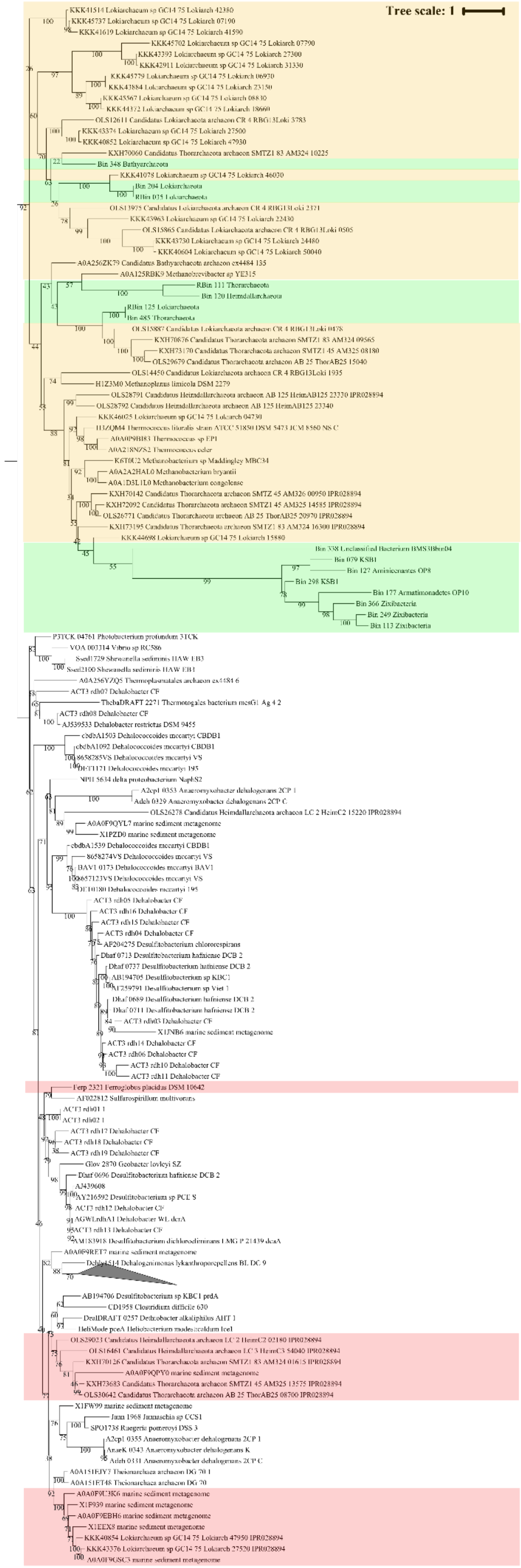
Tree scale: 1

**Additional file 12: Figure S11. Maximum-likelihood phylogenetic tree of putative dehalogenase in Shark Bay MDM MAGs.** Maximum-likelihood phylogenetic tree constructed with reductive dehalogenase domain (IPR028894) found in the MDM MAGs with 1000 bootstrap replications. Both reductive dehalogenase domain (IPR028894) and epoxyquiuosine reductase were found in Asgard archaea, KSB1, Aminicenantes (OP8), Armatimonadetes (OP10), Zixibacteria and Bathyarchaeota. Although these MAGs encode both epoxyquiuosine reductase and reductive dehalogenase domain, they cluster with homologous sequences of dehalogenase reductases. Thus it is unclear if the MDM community in Shark Bay can respire organohalides. Red shading indicates *bona fide* dehalogenases found in previous studies [6, 22], yellow shading indicates homologous sequences of dehalogenase reductases, and green shading represent reductive dehalogenase domains (IPR028894) in this study.

**Additional file 13: Table S1. Genome statistics of 24 high quality MDM MAGs and 91 medium quality MDM MAGs.**

**Additional file 14: Table S2. Rhodopsin sequences, BLAST results and reference rhodopsin sequences used in Additional file 6: Figure S5.**

**Additional file 15: Table S3. Table indicating the presence and absence of a wide range of genes involved in different metabolic pathways. Green boxes indicate presence of genes while white boxes indicate absence of genes.**

**Additional file 16: Table S4. RuBisCo sequences, BLAST results and reference RuBisCo sequences used in Fig. 5.**

**Additional file 17: Table S5. Relative abundance of the bacterial community in Shark Bay microbial mats. Green boxes indicate bacteria affiliated with microbial dark matter.**

**Additional file 18: Table S6. Relative abundance of the archaeal community in Shark Bay microbial mats. Green boxes indicate archaea affiliated with microbial dark matter.**

**Additional file 19: Table S7. Dissimilatory sulfate reduction sequences (*dsrAB*) identified in microbial dark matter MAGs in this study.**

**Additional file 20: Table S8. Reductive dehalogenase sequences identified in microbial dark matter MAGs in this study.**
